# Morphotype-specific calcium signaling in human microglia

**DOI:** 10.1101/2024.05.06.592754

**Authors:** Sofia Nevelchuk, Bianca Brawek, Niklas Schwarz, Ariel Valiente-Gabioud, Thomas V. Wuttke, Yury Kovalchuk, Henner Koch, Anke Höllig, Frederik Steiner, Katherine Figarella, Oliver Griesbeck, Olga Garaschuk

## Abstract

Key functions of Ca^2+^ signaling in rodent microglia include monitoring the brain state or the surrounding neuronal activity and sensing the danger or damage in their vicinity. Microglial Ca^2+^ dyshomeostasis is a disease hallmark in many mouse models of neurological disorders but the Ca^2+^ signal properties of human microglia remain unknown. Using a newly developed toolbox, we analyzed *in situ* Ca^2+^ signaling of decades-old human cortical microglia. The data revealed marked compartmentalization of Ca^2+^ signals, with signal properties differing across the compartments and resident morphotypes. The basal Ca^2+^ levels were low in ramified and high in ameboid microglia. The fraction of cells with ongoing Ca^2+^ signaling, the fraction and the amplitude of process Ca^2+^ signals and the duration of somatic Ca^2+^ signals decreased when moving from ramified via hypertrophic to ameboid microglia. In contrast, the size of active compartments, the fraction and amplitude of somatic Ca^2+^ signals and the duration of process Ca^2+^ signals increased along this pathway.

## Introduction

Microglia are the principal immune cells of the central nervous system (CNS), which are implicated in virtually all physiological (e.g., development, synaptic transmission, plasticity and sleep) and pathological (e.g., traumatic injury, glioma and neurodegenerative or autoimmune diseases) processes of the CNS (Garaschuk and Verkhratsky, 2019; Ma et al., 2024). A key aspect of microglial function is the ability to monitor their microenvironment and detect dyshomeostasis by sensing DAMPs (damage-) or PAMPs (pathogen-associated molecular patterns). For this purpose, they express a plethora of genes encoding different membrane receptors, the sum of which is referred to as microglial sensome (Hickman et al., 2013). In many cases, activation of those receptors leads to an increase in the intracellular free Ca^2+^ concentration ([Ca^2+^]i). Such transient changes in [Ca^2+^]i link the sensor and effector functions of microglia by triggering the generation/release of cytokines and other inflammatory factors (e.g., reactive oxygen species), proliferation, differentiation, migration and phagocytosis (Lushchak et al., 2021; Pan and Garaschuk, 2022). In mice, microglial Ca^2+^ transients are spatially compartmentalized and located in subcellular domains involved in a given (patho)physiological function. The normal function of cortical neural networks, for example, is accompanied by infrequent Ca^2+^ transients in microglial processes, with the frequency of process transients increasing dramatically during neural network hyper- or hypoactivity (Umpierre et al., 2020). Consistently, somatic Ca^2+^ transients are rare under homeostatic conditions but much more frequent during (neuro)inflammation, injury or damage (Brawek and Garaschuk, 2014; Brawek et al., 2017; Eichhoff et al., 2011; Pozner et al., 2015; Riester et al., 2020). Although under homeostatic conditions the mouse microglial sensome broadly resembles that of humans (Abels et al., 2021; Galatro et al., 2017; Gosselin et al., 2017), knowledge about the Ca^2+^ dynamics of human microglia is scarce and so far restricted to cultured primary microglia (e.g., Janks et al., 2018) or cultured human induced pluripotent stem cell (iPSC)-derived microglia-like cells (e.g., Granzotto et al., 2024; Jairaman et al., 2022; Jantti et al., 2022; Konttinen et al., 2019; Que et al., 2023).

Moreover, the morphological appearance of microglia in the intact human brain is indicative of a higher state of alertness compared to the mouse microglia (Franco Bocanegra et al., 2018; Torres-Platas et al., 2014). Whereas in mice the vast majority of cortical microglia, for example, show a ramified phenotype, microglia in the human cortex also appear in a primed, reactive, or amoeboid morphology. The dominance of ramified microglia over other morphotypes in mice is likely due to the specific pathogen-free environment of laboratory animals, which differs a lot from the typical environment of a human individual. In addition, the turnover rate of microglia in the human CNS is lower than that in the mouse CNS (Askew et al., 2017; Reu et al., 2017). This poses the question of whether human and mouse microglia differ in terms of their Ca^2+^ signaling.

To answer this question, we studied the Ca^2+^ signaling of human microglia residing in the spare cortical tissue obtained during glioblastoma/astrocytoma surgery or hippocampectomy (Table 1). The cells were lentivirally labeled with a newly developed ratiometric Ca^2+^ indicator mCyRFP1/CaNeon in human organotypic cortical slices cultured in human cerebrospinal fluid (hCSF). This preparation preserves well the cytoarchitecture and electrophysiological properties of excitatory and inhibitory neurons for several weeks (Schwarz et al., 2017; Schwarz et al., 2019).

**Table 1.**

| Age at surgery (y) | Sex | Resected brain area | Surgery cause |
| --- | --- | --- | --- |
| 56 | F | left temporal cortex | glioblastoma, WHO grade IV |
| 45 | F | right temporal cortex | diffuse astrocytoma, WHO grade II |
| 38 | F | right temporal cortex | astrocytoma, WHO grade II |
| 37 | M | right frontal cortex | astrocytoma, WHO grade III |
| 11 | F | right temporal cortex | hippocampal sclerosis right |
| 41 | M | right parietal cortex | intraventricular glioneuronal tumor (high grade) |

## Results

### New genetically encoded indicator for Ca^2+^ imaging in human microglia

To extend the methodology, previously developed for mice, to human tissue, we first infected the organotypic cortical slices (Fig. 1A) with the 1:1:1 mixture of miR9- regulated lentiviral vectors encoding mCherry (red=R), mVenus (green=G), and mTurquoise2 (blue=B) by adding 1-2 µl of the virus mixture (Fig. 1B) to the surface of brain slices during culturing. Because of the high endogenous activity of miR9 in most brain cells except microglia, this approach favors the specific labeling of microglial cells (Akerblom et al., 2013; Olmedillas et al., 2023). As shown in Fig. 1C, this protocol resulted in strong labeling of cells with microglia-like morphology. Due to the stochastic nature of viral transduction, individual cells expressed either one of the three fluorophores or any combination thereof (Fig. 1C and D).

**Figure 1.**
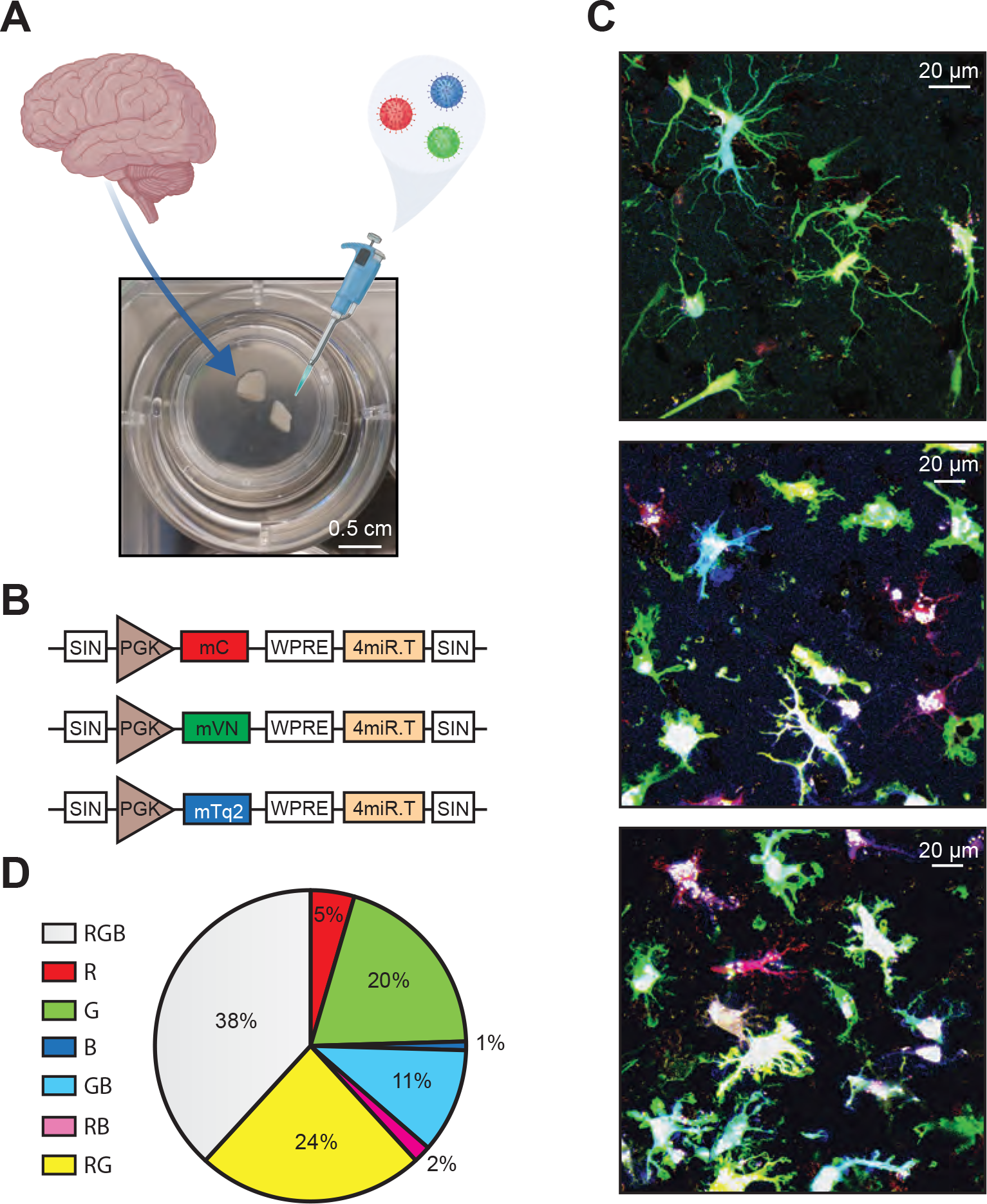
**miR-9-assisted labeling of human microglia**. (**A**) Schematics illustrating preparation, culturing and RGB labeling of human organotypic brain slices. (**B**) Scheme of the miR-9-regulated viral constructs inducing expression of mCherry (red), mVenus (green) and mTurquoise (blue) in microglia. (**C**) Maximum intensity projection (MIP) images (2-20 µm depth, here and below step size 1 µm) showing RGB-labeled human microglia in organotypic slices. (**D**) Pie chart showing the fractions of different colors in the RGB-labeled microglial population (n = 110 cells).

Next, we set out to develop an indicator suited for assessing the [Ca^2+^]i in human microglia. We opted for a ratiometric red/green indicator (Fig. 2A), which is less sensitive to movement artifacts and thus facilitates the identification of thin mobile processes, and engineered it to enable simultaneous fluorophore excitation in single- (excitation wavelength 488 nm) and two-photon modes (Fig. 2B).

**Figure 2.**
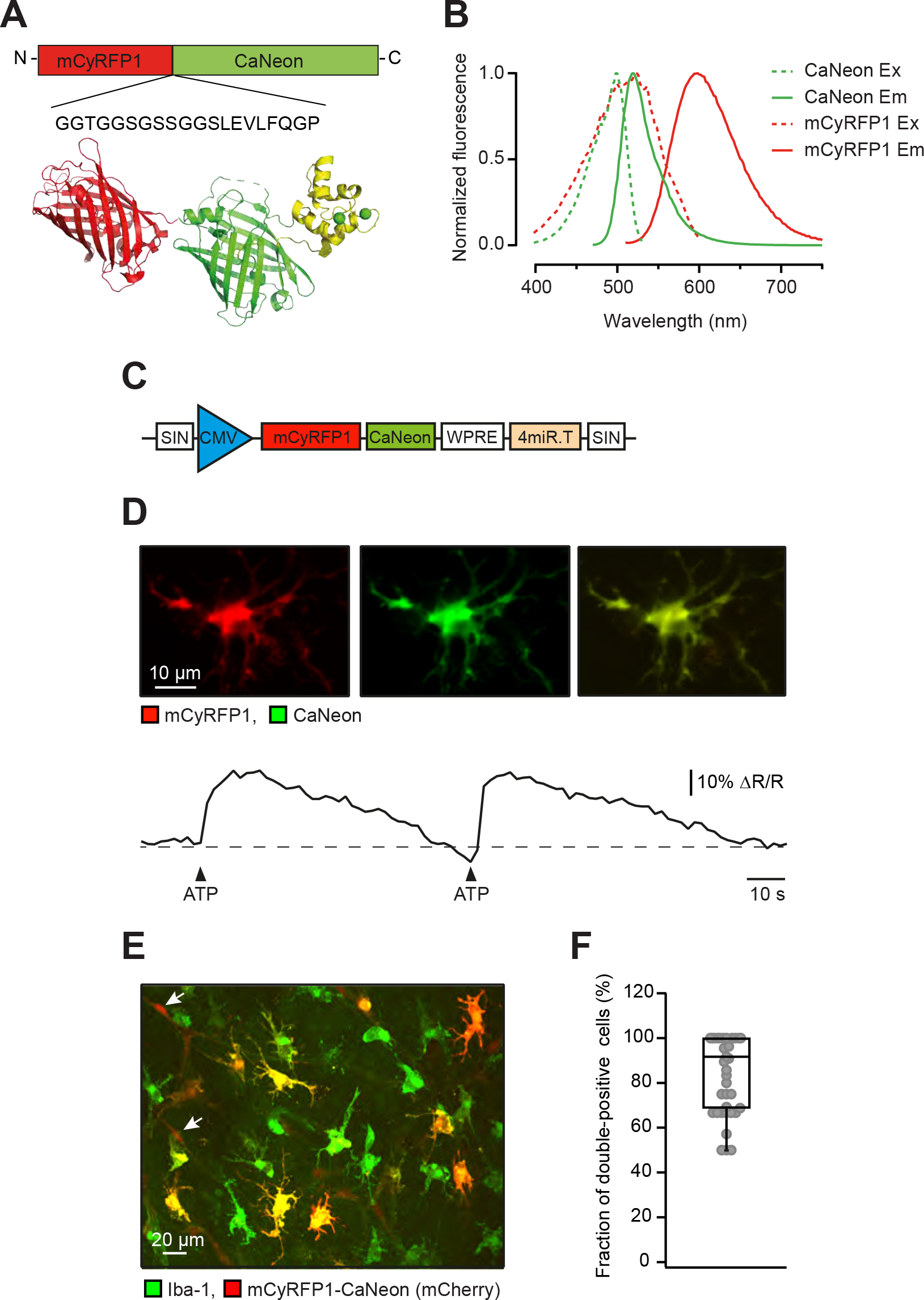
Functional properties of the new ratiometric Ca^2+^ sensor mCyRFP1- CaNeon. (**A**) Schematics of the sensor design (see Materials and Methods for details). (**B**) The excitation and emission spectra of purified mCyRFP1 (red) and CaNeon (green). Note that the excitation spectra show significant overlap, enabling the efficient one-photon excitation of the fluorophore by a single light source, and the emission spectra can be well separated. (**C**) Scheme of the miR-9-regulated viral construct for expressing mCyRFP1-CaNeon in *in situ* microglia. (**D**) Sample Ca^2+^ transients (lower panel) evoked by pressure application (12 psi, 200 ms) of 5 mM ATP (dissolved in the standard pipette solution: 150 mM NaCl, 2.5 mM KCl, 10 mM HEPES, pH 7.4) to a mCyRFP1-CaNeon-expressing cell (upper panel). (**E**) MIP (6-16 µm depth) image of a fixed organotypic slice, labeled with antibodies against a microglia/macrophage marker Iba-1 (green) and mCyRFP1-CaNeon (red). (**F**) Box plot showing the fraction of double- positive cells among the cells, positive for mCyRFP1-CaNeon (n = 254 mCyRFP1- CaNeon -positive cells, 41 FOV).

Thus, we first constructed CaNeon, a non-ratiometric Ca^2+^ indicator consisting of the yellow-green fluorescent protein mNeonGreen (Shaner et al., 2013) and a minimal Ca^2+^ binding domain derived from Troponin C (TnCmin) with only 2 Ca^2+^ binding sites (versus 4 in GCaMPs), thus lowering the buffer capacity of the indicator (Fig. S1A-D). We started from GreenT-EC, which is tuned to monitor Ca^2+^ levels in interstitial fluids (Valiente-Gabioud et al., 2023), and derived a variant that binds Ca^2+^ with high affinity while retaining large fractional fluorescence changes (Fig. S1 and Fig. 2). We then conjugated this indicator to the red fluorescent protein mCyRFP1 (Laviv et al., 2016) (Fig. 2A). For benchmarking, the spectroscopic *in vitro* properties of CaNeon and mCyRFP1-CaNeon were compared to the established Ca^2+^ sensor GCaMP6f (Chen et al., 2013). The extinction coefficient, quantum yield and Ca^2+^ sensitivity of both indicators (Fig. S1D) were similar to that of GCaMP6f but with better linearity (Hill coefficient 1.5-1.6 instead of 2.3).

Taking into account our previous experience in mice (Brawek et al., 2017), when preparing miR9-regulated lentiviral vectors encoding mCyRFP1-CaNeon we switched to a stronger promotor (CMV instead of PGK; Fig. 2C vs. Fig. 1B). From such a vector, the sensor expressed well both in Human Embryonic Kidney (HEK) 293 cells in culture (Fig. S1E-H) and human cortical microglia in organotypic slices (Fig. 2D-F). For initial testing of the new construct, HEK 293 cells were stimulated by a bath application of caffeine (Fig. S1F-H), an agonist of ryanodine receptors releasing Ca^2+^ from the intracellular Ca^2+^ stores (Garaschuk et al., 1997), and microglia – by a local pressure application (250 ms, 12 psi) of the purinergic receptor agonist ATP (Fig. 2D), resulting in both cases in large Ca^2+^ transients.

The specificity of the miR9-regulated microglial labeling was assessed in fixed organotypic slices, labeled with antibodies against a microglia/macrophage-specific marker Ionized Ca^2+^-binding adaptor molecule 1 (Iba-1) and mCherry, which has 63% and 33% homology with mCyRFP1 and CaNeon, respectively (Fig. 2E). In fixed tissue, the native fluorescence of CaNeon was too low to be visible in the green channel. While 91.67 ± 32.29% (Median ± IQR; 254 cells/41 fields of view (FOVs)) of mCyRFP1- CaNeon-positive cells were also Iba-1-positive (Fig. 2F), in the human brain we did identify another weekly-labeled Iba-1-negative population of spindle-shaped cells, often associated with blood vessels (presumed pericytes, arrows in Fig. 2E and Fig. S2). The latter, however, were dim and therefore hardly visible *in situ* and were morphologically too different to be mistaken for microglia. These cells were excluded from further analyses.

Taken together, the novel technique described above enables real-time morphological and functional analyses of human microglia, including longitudinal analyses of process/cell body motility (Movie S1) as well as Ca^2+^ imaging.

### Morphotype-specific properties of microglial Ca^2+^ signals

In contrast to the mouse cerebral cortex, in which under homeostatic conditions more than 90% of microglia have ramified morphology, in the human postmortem cortex microglial cells appear much more heterogeneous, with only half of the population showing ramified morphology and another half being composed of hypertrophic (also called reactive; 32%) and ameboid (18%) cells (Torres-Platas et al., 2014). Consistently, out of 81 cells analyzed in this study, 46.91% were ramified, 30.86% hypertrophic and 22.22% ameboid (Fig. 3A). All 3 microglial morphotypes were also present in freshly resected human cortical tissue (Fig. S3). However, our *in situ* microglia were larger. Some 20% of them had a mean diameter going beyond that of cells from freshly resected tissue (Fig. S3B), whereas the distributions of soma shapes looked similar (Fig. S3E).

**Figure 3.**
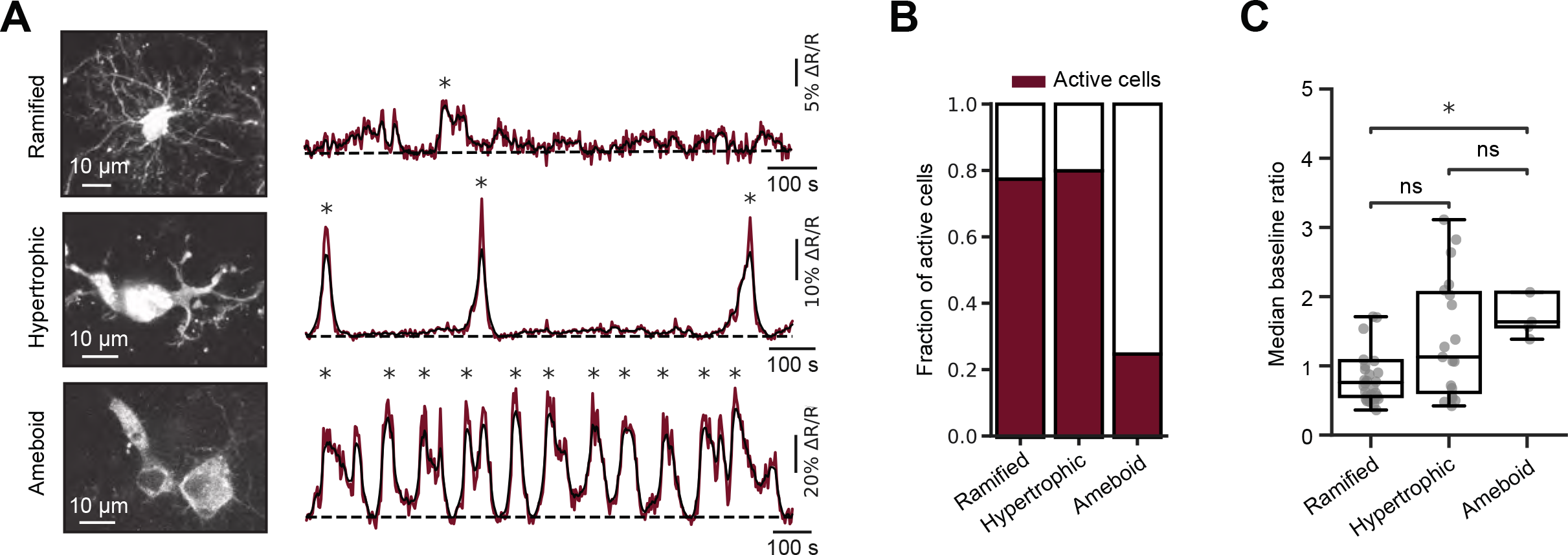
Ongoing Ca^2+^ signaling in microglia of different morphology. (**A**) MIP images showing examples of mCyRFP1-CaNeon-positive microglia with ramified, hypertrophic, and amoeboid morphology (left; 8-32 µm, 7-30 µm, and 3-16 µm depth, respectively) as well as whole-cell spontaneous Ca^2+^ signals (asterisks), recorded from these cells during a 15-min-long recording period (right). Dark red, original traces, black, traces filtered with Gaussian filter ensemble (σ = [0.5, 1, 2, 3, 4] \ 5), (**B**) Bar graph summarizing the fractions of active cells of different morphologies (n=38 ramified, 25 hypertrophic and 18 ameboid microglia). (**C**) Box plot illustrating the distributions of the median (per cell) basal ratios of ramified, hypertrophic and ameboid microglia (P=1.2*10^-2^ for comparison between ramified and ameboid microglia, Kruskal-Wallis test; n=31, 20, 5 active cells, respectively).

Different morphotypes also differed in terms of their Ca^2+^ signaling. Like their mouse counterparts *in vivo* (Eichhoff et al., 2011, Umpierre et al., 2020), human microglia in organotypic slices showed ongoing Ca^2+^ signaling (Fig. 3A). While the fractions of active cells (showing at least one Ca^2+^ transient at any subcellular location during the 15-min-long recording period) were approximately similar among ramified and hypertrophic microglia (Fig. 3B), the patterns of Ca^2+^ transients sampled over the entire cell looked different, with an increase in the frequency and amplitude of these transients from ramified to ameboid microglia (Fig. 3A) and a significant reduction in the fraction of active cells for ameboid microglia (Fig. 3B; P=3*10^-4^, Chi-squared test). Moreover, the median basal ratios (proportional to the basal [Ca^2+^]i) of active cells increased gradually from ramified through hypertrophic to ameboid cells, with the difference between the ramified and ameboid cells reaching the level of statistical significance (Fig. 3C; P=1.2*10^-2^, Kruskal-Wallis test). In 3 hypertrophic and 1 ameboid cell, ongoing Ca^2+^ transients showed oscillatory behavior (e.g., Fig. 3A, lower right panel) with a median frequency of 12.33*10^-3^ s^-1^. In 4 recorded cells the microglial processes formed structures, reminiscent of phagocytic cups, with cups often showing compartmentalized Ca^2+^ transients (Movie S2). Although we did not observe any somatic movements over a 15-minute-long recording period, many ramified and hypertrophic cells showed vivid process motility, also often associated with compartmentalized Ca^2+^ transients (Movie S1).

Therefore, the subsequent analyses of Ca^2+^ signaling were conducted based on active subcellular compartments (regions of activity, ROA) using the Begonia ROA detection algorithm (Bjornstad et al., 2021; see Materials and methods for details). On average, individual microglial cells contained 2.53 ± 0.35 (mean ± SEM; n= 56 cells) active subcellular compartments, some of which are exemplified in Fig. 4A. Out of 78 ROA detected in ramified cells, 75.6% were localized to the processes, 11.5% covered soma and processes and 12.8% covered cell somata only (Fig. 4B). Surprisingly, we have rarely observed global Ca^2+^ signals invading the entire cell. The spatial distribution of Ca^2+^ signals in hypertrophic cells was similar to that in ramified microglia (P=0.31, Chi-squared test). In ameboid cells, however, the spatial pattern of Ca^2+^ signals was entirely different, with 66.7% of ROA localized to cell somata (P<10^-3^, Chi-squared test). Consistently, the areas of individual ROAs were small in ramified, somewhat larger and heterogeneous in hypertrophic and significantly larger in ameboid cells (P=2.3*10^-3^, Kruskal-Wallis test; Fig. 4C). The cell type-specific frequency of ongoing Ca^2+^ signals tendentially showed a bell-shaped form, with the highest median frequency observed in hypertrophic cells (Fig. 4D). The differences, however, did not reach the level of statistical significance. Next, we characterized the properties of Ca^2+^ transients derived from different ROA in the 3 morphotypes (Fig. 4E-H). The median amplitudes of Ca^2+^ transients were significantly smaller in hypertrophic, compared to ramified and ameboid, microglia (Fig. 4F). The durations (full width at half maximum, FWHM) were larger and the total Ca^2+^ load (area under the curve, AUC) was smaller in hypertrophic compared to ramified cells (Fig. 4, G and H).

**Figure 4.**
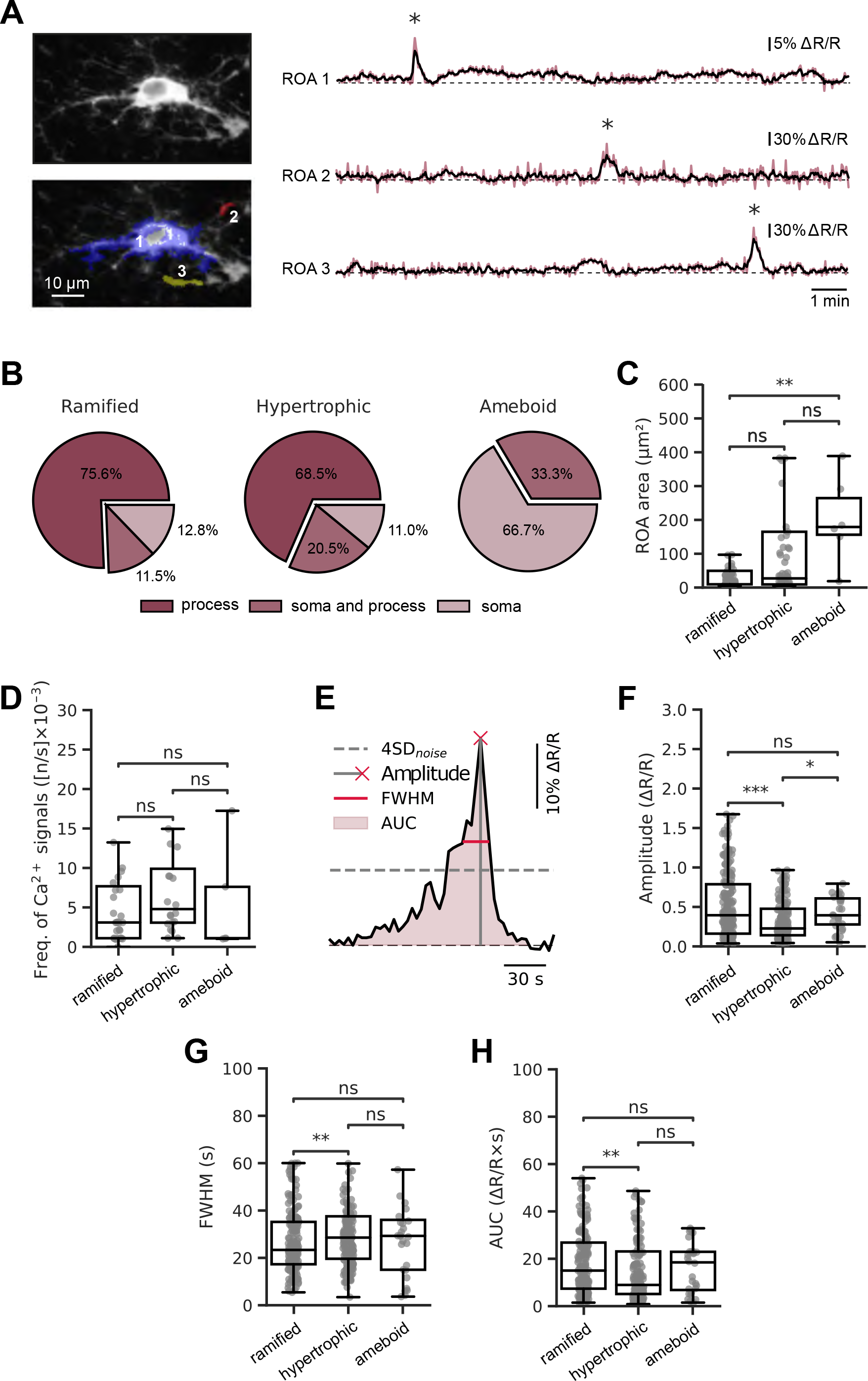
Subcellular compartmentalization of ROAs in human microglia. (**A**) Left: 4D average intensity projection of a mCyRFP1-CaNeon-expressing ramified microglial cell alone (upper panel) and with the overlayed sample ROAs, shown in different colors (lower panel). Right: spontaneous ongoing Ca^2+^ signals (asterisks) recorded from ROAs, labeled with the respective number in the lower left panel. (**B**) Pie charts showing the spatial distribution of ROA in ramified (left panel), hypertrophic (middle panel), and amoeboid (right panel) microglia. (**C**) Box plots showing the morphotype- specific distributions of ROA areas. ROAs were significantly smaller in ramified compared to amoeboid microglia (P=10^-3^; here and below: the Kruskal-Wallis test followed by the Holm-Bonferroni post hoc test for multiple comparisons). (**D**) Box plots illustrating the frequencies per cell of Ca^2+^ transients in ramified, hypertrophic and ameboid microglia (n=31, 20, 5 cells, respectively). (**E**) Schematic, defining the parameters of Ca^2+^ transients analyzed in this study. (**F-H**) Box plots showing amplitudes (**F**; P=3.4*10^-4^ and 0.02 for comparison of ramified to hypertrophic and hypertrophic to ameboid microglia, respectively), FWHM (**G**; P=8.7*10^-3^ for comparison of ramified to hypertrophic microglia) and AUC (**H**; P=3.3*10^-3^ for comparison of ramified to hypertrophic microglia) of Ca^2+^ transients for microglia of different morphologies.

Detailed analyses of individual Ca^2+^ transients revealed that although the majority of ROAs in ameboid cells were located in cell somata (Fig. 4B), the soma and process ROAs showed many more Ca^2+^ transients (Fig. 5A). Moreover, for ramified and hypertrophic morphotypes process Ca^2+^ transients had the highest amplitudes and AUCs, whereas their durations varied between the morphotypes (Fig. S4). In ramified microglia, the durations of process Ca^2+^ transients were the shortest (Fig. S4B), whereas in hypertrophic and ameboid microglia the durations of Ca^2+^ transients localized to soma and processes became gradually longer than that of somatic Ca^2+^ transients, reaching the level of significance in ameboid microglia (Fig. S4E and H). The data also revealed the compartment-specific heterogeneity of Ca^2+^ transients in ramified, hypertrophic and ameboid microglia and prompted the compartment-specific comparison across morphotypes (Fig. 5B-J). In general, somatic Ca^2+^ transients of ameboid cells had the largest amplitude and the shortest duration (Fig. 5B and C), whereas ramified cells had the largest AUCs (Fig. 5D). For ROA located in soma and processes, the amplitudes and AUCs increased gradually from ramified through hypertrophic to ameboid cells (Fig. 5E and G). The process Ca^2+^ transients had higher amplitudes and shorter durations in ramified microglia (Fig. 5H and I).

**Figure 5.**
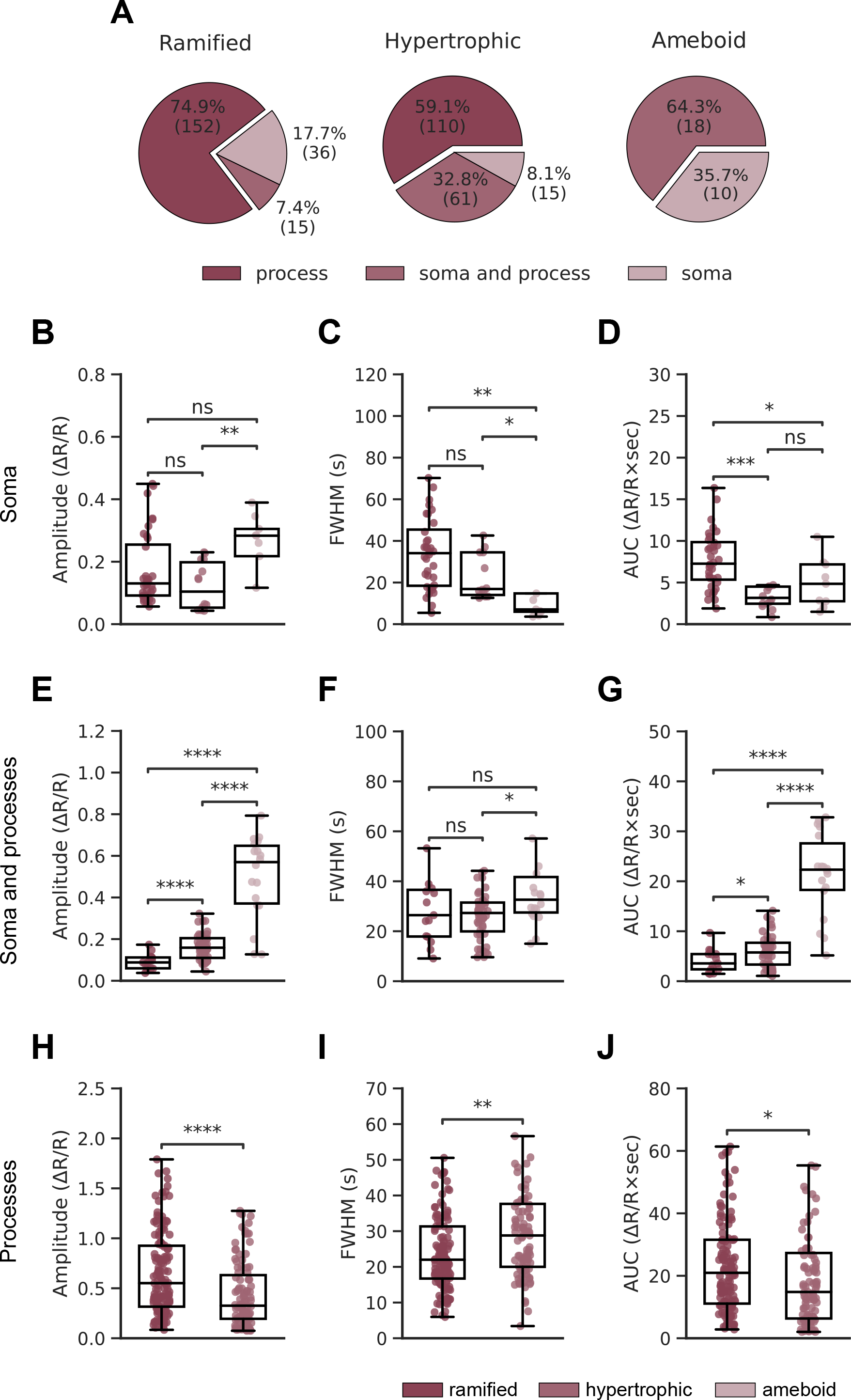
Characteristics of microglial Ca^2+^ transients in different subcompartments. (**A**) Pie charts showing the distribution of Ca^2+^ transients among different subcompartments in ramified (left panel), hypertrophic (middle panel), and amoeboid (right panel) human microglia. (**B**-**D**) Box plots comparing amplitude (**B**; P=6*10^-3^ for comparison of hypertrophic to ameboid microglia (here and below: the Kruskal-Wallis test followed by the Holm-Bonferroni post hoc test for multiple comparisons)), FWHM (**C**; 1.4*10^-3^ and 0.01 for comparison of ramified to ameboid and hypertrophic to ameboid microglia, respectively), and AUC (**D**; P=0.03 and 2.2*10^-4^ for comparison of ramified to ameboid and ramified to hypertrophic microglia, respectively) of somatic Ca^2+^ transients in ramified, hypertrophic and amoeboid microglia. (**E-G**) Box plots comparing amplitude (**E**; P=4.8*10^-5^, 2.1*10^-6^ and 4.6*10^-7^ for comparison of ramified to hypertrophic, ramified to ameboid and hypertrophic to ameboid microglia, respectively), FWHM (**F**; P=0.01 for comparison of hypertrophic to ameboid microglia), and AUC (**G**; P=3.7*10^-2^, 4.4*10^-6^ and 1.8*10^-8^ for comparison of ramified to hypertrophic, ramified to ameboid and hypertrophic to ameboid microglia, respectively) of Ca^2+^ transients in soma and processes of ramified, hypertrophic and amoeboid microglia. (**H-J**) Box plots comparing amplitude (**H**; P=3.1*10^-5^), FWHM (**I**; P=1.3*10^-3^), and AUC (**J**; P=0.02) of process Ca^2+^ transients of ramified and hypertrophic microglia.

Thus, our data revealed the marked compartmentalization of Ca^2+^ transients in human microglia. Interestingly, the basic properties of these Ca^2+^ transients differed dramatically across the compartments as well as morphotypes.

## Discussion

In this study, we introduced a new approach for functional analyses of *in situ* human microglia cultured in hCSF and applied it to characterize the ongoing Ca^2+^ signaling in microglial morphotypes (i.e., ramified, hypertrophic and ameboid microglia), present in the human brain. Our data revealed distinct functional properties of each morphotype. Whereas in ramified microglia basal [Ca^2+^]i was low and ongoing Ca^2+^ transients were mostly compartmentalized in microglial processes, having large amplitudes and short durations, the ameboid microglia had significantly higher basal [Ca^2+^]i and the vast majority of large amplitude Ca^2+^ signals invaded cell somata. Moreover, the fraction of cells with ongoing Ca^2+^ signaling, the fraction and the amplitude of process Ca^2+^ signals and the duration of somatic Ca^2+^ signals decreased when moving along the microglia activation pathway, i.e. from ramified via hypertrophic to ameboid microglia (Fig. 6). In contrast, the size of active compartments, the fraction and the amplitude of somatic Ca^2+^ signals and the duration of process Ca^2+^ signals increased with microglial activation (Fig. 6). While imaged microglial cells often showed overt process movement associated with localized Ca^2+^ signals, within the framework of the present study (15- min-long recording period) we have not observed any translocations of cell somata.

**Figure 6.**
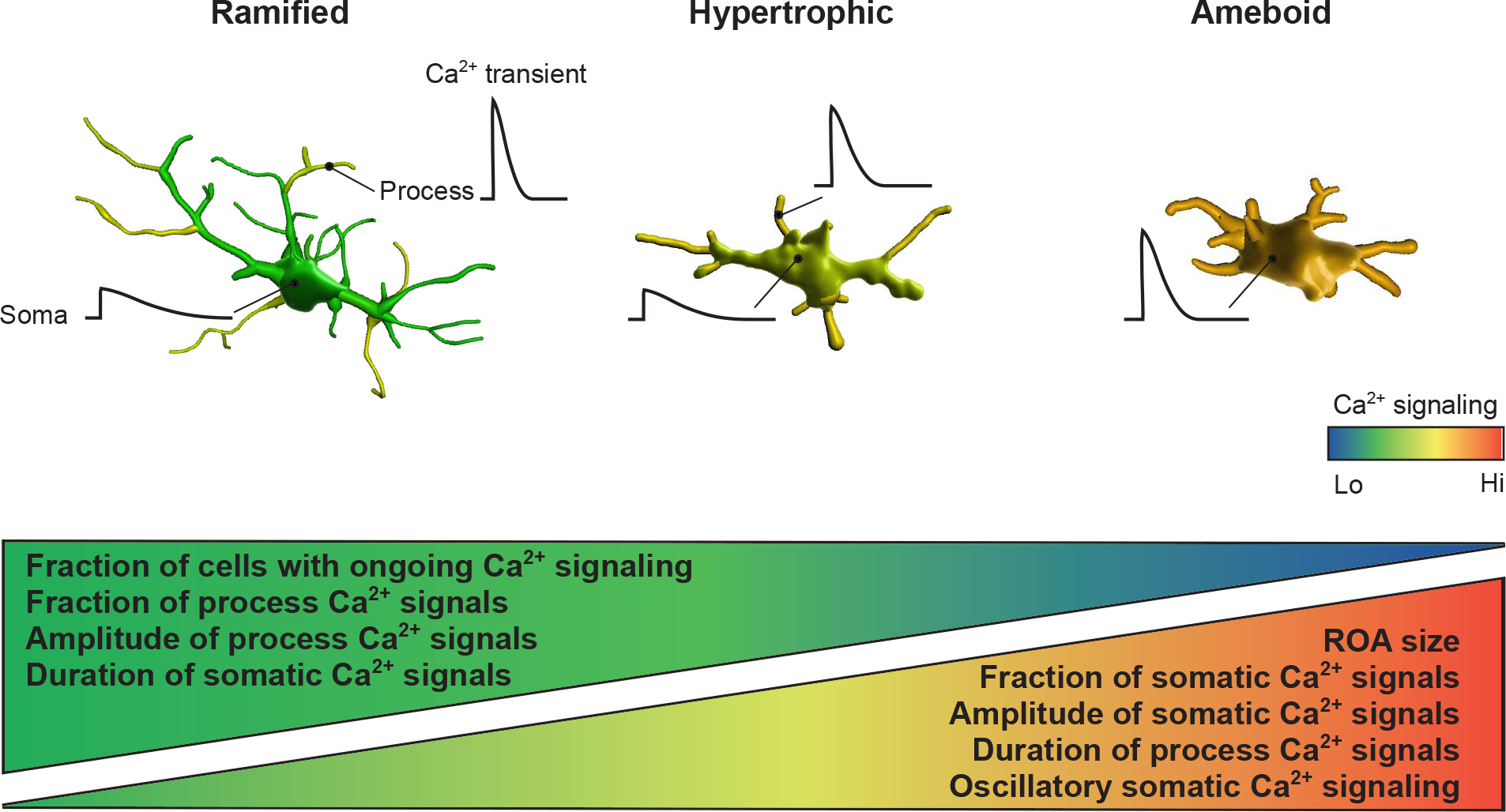
Ongoing Ca^2+^ signals in human microglia are subcompartment- and morphotype-specific. Graphical summary of main findings. See discussion for details.

To enable Ca^2+^ imaging of human microglia, we combined several cutting-edge techniques, recently developed by our laboratories. Thus, organotypic brain slices derived from fresh human biopsies were cultured in human CSF. This approach preserves well not only neurons (Schwarz et al., 2017; Schwarz et al., 2019) but also different morphotypes of human microglia (Fig. 1). Contrary to *in vivo* mouse cortex with or without transplanted human cortical organoids, which contains ramified human or mouse microglia (Nimmerjahn et al., 2005, Mancuso, 2019 #244; Schafer et al., 2023), and cultured human iPSC-derived microglial cells, which are largely ameboid (Granzotto et al., 2024; Jairaman et al., 2022; Jantti et al., 2022; Konttinen et al., 2019; Que et al., 2023), our model harbors all morphotypes, thus allowing their comparative study. Moreover, throughout the experiment, the microglia dwell in their native microenvironment. This is important because of the well-known environment-induced cell-non-autonomous shifts in microglial phenotype (Gosselin et al., 2017; Schafer et al., 2023). The distribution of ramified, hypertrophic and ameboid microglia in our preparation was close to that found in fresh-frozen postmortem cortical samples (Franco Bocanegra et al., 2018; Torres-Platas et al., 2014). Accordingly, we have also found all 3 morphotypes in acutely resected human cortical tissue. The soma size of these cells, however, was smaller than that of *in situ* microglia. While some inter-individual as well as surgery-related variability cannot be excluded, this difference is likely caused by organotypic slice culturing.

The use of RGB labeling provides each microglial cell with a unique color tag and enables the longitudinal monitoring of its behavior, including cell death or division, phagocytosis, somatic dislocation or process movement under control conditions and, in the future, also in human disease models (McGeachan et al., 2024). Finally, the development of a novel genetically encoded Ca^2+^ indicator mCyRFP1-CaNeon, well-suited for single- as well as two-photon-based ratiometric Ca^2+^ imaging, adds to the versatile microglia analyses toolbox and has enabled us to measure the morphotype-specific basal Ca^2+^ levels.

Careful identification of all active pixels within the cell (Fig. 4), allowed us to analyze the spatial dimension of microglial Ca^2+^ signaling. The data revealed that microglial Ca^2+^ signals are highly compartmentalized and rarely spread over the entire cell. Moreover, in ramified microglia, the amplitudes and time courses of Ca^2+^ transients differed significantly between different subcompartments (Fig. 6), with large and short process Ca^2+^ transients likely targeting different signaling pathways compared to small but prolonged somatic Ca^2+^ signals (Dolmetsch et al., 1998; Wang et al., 2023).

This study uniquely analyzed Ca^2+^ signaling in microglia stemming from decades-old brains, in sharp contrast to recent analyses of iPSC-derived microglia, generated, on average, 30-50 days before the experiment (Granzotto et al., 2024; Jairaman et al., 2022; Que et al., 2023). Consistently, the properties of the two cell populations differed substantially both in morphology (largely ameboid iPSC-derived microglia vs. 3 different morphotypes analyzed in our study) and function. While iPSC-derived microglia moved vividly around the dish with the mean speed of 5 µm/min (Granzotto et al., 2024; Jairaman et al., 2022), we did not observe any translocation of microglial somata over the 15-min-long recording period, reminiscent to *in vivo* data from adult mice (Bolmont et al., 2008; Mondo et al., 2020; Olmedillas et al., 2023). Consistent with their morphology and similar to our ameboid cells, iPSC-derived microglia showed global Ca^2+^ signals invading both somata and processes (Que et al., 2023), quite in contrast to much more compartmentalized Ca^2+^ signaling patterns of ramified and hypertrophic microglia studied here. It also has to be kept in mind that in contrast to iPSCs, microglia stem from the extra-embryonic yolk sack macrophages (Ginhoux et al., 2010).

So far, Ca^2+^ signaling of ramified microglia was exclusively studied in mice under *in situ* (Logiacco et al., 2021; Seifert et al., 2011) or *in vivo* (Brawek and Garaschuk, 2014; Brawek et al., 2017; Eichhoff et al., 2011; Pozner et al., 2015; Umpierre et al., 2020; Xia et al., 2021) conditions. However, recent comparative mouse-human studies revealed several functional differences between the human and mouse microglia. First, human microglia renew at a median rate of 28% per year (Reu et al., 2017), which means that some microglia dwell in the human brain for more than two decades, whereas mice live ∼ 2 years and the oldest mouse microglia in which Ca^2+^ signaling was studied were 18-21-month-old (Olmedillas del Moral et al., 2020). Secondly, despite the overall similarity between the genes expressed by human and mouse microglia, several immune genes do not have a clear 1:1 mouse ortholog (e.g., complement receptor type 1, HLA-DRB1, HLA-DRB5, TLR, Fcγ and SIGLEC receptors) or display <60% identity between human and mouse at the primary amino acid sequence (e.g. TREM2), and several rodent gene modules, including complement, phagocytic and neurodegeneration-susceptibility genes, differed substantially from primate microglia (Galatro et al., 2017; Geirsdottir et al., 2019; Mancuso et al., 2019). Moreover, whereas in most species microglia showed a single dominant transcriptional state, human microglia displayed substantial heterogeneity (Geirsdottir et al., 2019) and, for example, responded to neurodegeneration and Alzheimer’s disease pathology by upregulation of TMEM119, P2RY12 and CX3CR1 genes, contrary to their downregulation in mouse models of the disease (Keren-Shaul et al., 2017; Zhou et al., 2020).

Against this backdrop, we have compared the ongoing Ca^2+^ signaling in human and mouse microglia and have spotted several major similarities. First, the studied human cells seemed to belong to a continuum of microglial functional states with ramified and ameboid microglia building the two extremes and hypertrophic microglia spreading among multiple intermediate states (Hanisch and Kettenmann, 2007). Moreover, (i) the basal [Ca^2+^]i increased with the degree of activation both in human (Fig. 3C) and mouse (Brawek et al., 2017) microglia; (ii) both species showed spontaneous ongoing Ca^2+^ signaling; (iii) microglial Ca^2+^ transients in both species were highly compartmentalized (Fig. 4; Pan and Garaschuk, 2022; Umpierre et al., 2020) and (iv) the spatial distribution of ongoing Ca^2+^ signals switched from process- to soma-dominated patterns during microglial activation (Figs. 3-5; Eichhoff et al., 2011; Pozner et al., 2015; Umpierre et al., 2020).

Collectively, our data establish biopsy-derived organotypic human brain slices, cultured in human CSF, as a valid model for functional analyses of human microglia. This model provides a unique possibility to study adult, decades-old microglia in their natural microenvironment and reveals that many functional properties of these cells, including vivid process motility and compartmentalized ongoing Ca^2+^ signaling, the pattern of which is morphotype-specific, are reminiscent of that seen *in vivo* in rodent microglia.

## Acknowledgments

We thank E. Zirdum, B. Gittel and K. Schmidt for technical assistance, Radu-Gabriel Copie and Friederike Pfeiffer for the data shown in Fig. S2B, and Kuang Pan for the help with data acquisition. This work was partially supported by the DFG, grant number GA 654/13-2 (FOR2715) to O.G. The authors declare no competing financial interests. Author contributions: S. Nevelchuk: data analysis, visualization; B. Brawek: visualization, writing; N. Schwarz: slice preparation and culturing, provided critical reagents, writing; A. Valiente-Gabioud: generation of the Ca^2+^ sensor; T.V. Wuttke: tissue resection, provided critical reagents; Y. Kovalchuk: investigation; H. Koch: provided critical reagents; A. Höllig: tissue resection; F. Steiner: investigation, cloning; K. Figarella: prepared all viral vectors; O. Griesbeck: experimental design, generation of Ca^2+^ sensor, writing; Olga Garaschuk: conceptualization, experimental design, investigation, supervision, visualization, and writing.

## Materials and methods

### Tissue specimen

Organotypic cortical slices were prepared from the cortical tissue, surgically resected to gain access to the pathology (Table 1). Approval (# 772/2021BO2) of the ethics committee of the University of Tübingen as well as written informed consent was obtained from all patients, whose resected tissue was used in this study.

Immunohistochemical data shown in Fig. S3 were obtained from the cortical tissue, surgically resected to gain access to the pathology and fixed immediately after resection (Table 2). Approval (EK067/20) of the ethics committee of the RWTH Aachen University as well as written informed consent for tissue donation and approval of the use of human tissue for scientific purposes and all related experimental procedures was collected before the commencement of the study.

**Table 2.**

| Age at surgery (y) | Sex | Resected brain area | Surgery cause |
| --- | --- | --- | --- |
| 74 | F | right frontal cortex | Metastasis (non-small cell lung cancer) |
| 34 | M | left orbitofrontal cortex | Diffuse glioneuronal tumor with oligodendroglioma-like features and nuclear clusters |
| 75 | F | right frontal cortex | Glioblastoma |
| 73 | M | left temporal cortex | Glioblastoma |

### Slice preparation and culturing

Slice preparation and culturing were performed as detailed previously (Schwarz et al., 2017; Schwarz et al., 2019). Briefly, the cortex was resected, microdissected and immediately transferred into the ice-cold artificial cerebrospinal fluid (aCSF) of the following composition (in mM): 110 choline chloride, 26 NaHCO3, 10 D-glucose, 11.6 Na-ascorbate, 7 MgCl2, 3.1 Na-pyruvate, 2.5 KCl, 1.25 NaH2PO4, und 0.5 CaCl2, pH 7.4, when bubbled with 95% O2, 5% CO2. The tissue was kept submerged in cool and oxygenated aCSF at all times. Slices (thickness 250-350 µm) were cut using a vibratome (Microm HM 650V, Thermo Fisher Scientific Inc, USA) and transferred onto culture membranes for cultivation. For the first hour following the slicing procedure, the slices were cultured in 1.5 ml neural stem cell media (48% DMEM/F-12 (Life Technologies), 48% Neurobasal (Life Technologies), 1x N-2 (Capricorn Scientific), 1x B-27 (Capricorn Scientific), 1x Glutamax (Life Technologies), 1x NEAA (Life Technologies), 20 mM HEPES) before changing to 1.5 ml hCSF per well without any supplements. For transduction with lentiviral vectors, encoding either a combination of 3 different fluorophores (mCherry, mVenus, mTurquoise) or the novel genetically encoded Ca^2+^ indicator (GECI) mCyRFP1-CaNeon (see below), 1-2 µl of the respective virus or virus mixture was added to the surface of each section within the first 24 hours after resection. Thereafter, the slices were cultured for 4-6 days onto culture membranes (uncoated 30 mm Millicell-CM tissue culture inserts with 0.4 mm pores, Merck Millipore, Germany) and kept in six-well dishes (BD Biosciences) in 1.5 ml of hCSF, which was replaced every 3 days. The dishes were kept in an incubator (Thermo Scientific) and maintained at 37°C, 5% CO2 and 100% humidity.

### Novel Ca^2+^ indicator CaNeon

When developing the ultralow affinity Ca^2+^ sensor GreenT-EC (Valiente-Gabioud et al., 2023), we also obtained the crystal structure of an intermediate variant named NRS 1.2 (PDB 8COT). Starting from NRS 1.2 we developed CaNeon via several iterative rounds of optimizations and screenings. Compared to NRS1.2, CaNeon incorporates the following additional mutations: C159Y, D198G, G248D, S255C, I258R. These amino acid changes improved the Ca^2+^ binding properties of the sensor, as well as its expression in mammalian cells. The full protein sequence of CaNeon is displayed in Fig. S1A. For designing a ratiometric version of the sensor, we used the red fluorescent protein mCyRFP1 as a reference fluorophore. The spectroscopic properties of mCyRFP1 allow simultaneous one- and two-photon-based excitation of both mCyRFP1 and CaNeon with an efficient spectral separation of the emission channels (Laviv et al., 2016). The mCyRFP1 was cloned at the N-terminus of CaNeon using a 20 amino acid-long flexible hydrophobic linker (GGTGGSGSSGGSLEVLFQGP, Fig. 2A). All cloning steps were done using the homology-based SliCE method (Zhang et al., 2012). In the absence of Ca^2+^, the fluorescence of CaNeon is low (Fig. S1B). The binding of Ca^2+^ ions causes conformational changes in CaNeon, resulting in a strong increase in fluorescence. The spectroscopic *in vitro* properties of CaNeon and the fusion protein mCyRFP1-CaNeon are summarized in Fig. S1D.

### Protein purification and biophysical characterization of CaNeon

His-tagged proteins were expressed in E.coli BL21 (Invitrogen) overnight at 37 °C in 50 mL auto-inductive Luria-Bertani medium, supplemented with 0.05% D-(+)- glucose (w/v), 0.2% lactose (w/v), 0.6% glycerol (v/v). Bacteria were harvested by centrifugation (4 °C, 10 min, 6000 g) and re-suspended in 10 mL resuspension buffer (20 mM Na2PO4, 300 mM NaCl, 20 mM imidazole; Sigma Aldrich) supplemented with protease inhibitors (4 μM PMSF, 20 μg/mL Pepstatin A, 4 μg/mL Leupeptin; Sigma Aldrich), 5 μg/mL DNase and 10 μg/mL RNase (Sigma Aldrich). Resuspended bacteria were lysed through sonication on ice for 7 min (80% of the time on; Bandelin Sonoplus). Insoluble components were removed through centrifugation (4 °C, 30 min at 20000 g). The supernatant was incubated with 150 μL 6% (v/v) Nickel-IDA agarose bead suspension (Jena Bioscience) for 1 hour at 4 °C under mild agitation. Agarose beads were collected in 1 mL propylene gravity flow columns (Qiagen) and washed with 10 mL resuspension buffer. The proteins were collected using 800 μL elution buffer (20 mM Na2PO4, 300 mM NaCl, 300 mM imidazole; Sigma Aldrich) and dialyzed against MOPS buffer (30 mM MOPS (3-morpholinopropane-1-sulfonic acid), 100 mM KCl, pH 7.2) for further measurements.

The ratio change (ΔR/R0) of purified mCyRFP1-CaNeon was determined by measuring the fluorescence at 520 and 650 nm upon excitation at 488 nm in MOPS buffer supplemented with 10 mM EGTA or 0.2 mM Ca^2+^. The molar extinction coefficients (EC) were determined via quantifying protein concentrations using the absorption of the denatured chromophore at 452 nm (with an EC of 44 mM-1cm-1). Proteins were prepared in MOPS buffer supplemented with 0.2 mM CaCl2 and the absorbance spectrum was acquired before and after the addition of NaOH to a final concentration of 0.1 M. The quantum yield of CaNeon in mCyRFP1-CaNeon was determined relative to mNeonGreen using the slope method, measuring the absorbance and emission spectra of serial dilution of the proteins. For the quantum yield and extinction coefficient measurements, the formation of CyRFP1 chromophore was prevented by incorporating the mutation Y68C. This allowed us to determine the structural effect of the N-terminal tag on CaNeon without the spectral interference of mCyRFP1. The brightness was calculated as extinction coefficient ˣ quantum yield.

The Ca^2+^ affinity of the sensors was determined using MOPS buffer supplemented with 10 mM EGTA and 1 mM Mg^2+^ by increasing concentrations of Ca^2+^ as previously described (Tsien, 1989; Zhang et al., 2012). The dissociation constant (Kd) values were determined by plotting the log10 values of the free Ca^2+^ concentrations against the corresponding ΔF/F0 or ΔR/R0 values (normalized to the response at 39.8 μM Ca^2+^) and fitting a sigmoidal curve to the plot. The kinetic rates of the Ca^2+^ indicators were measured in a Varian Cary Eclipse fluorescence spectrophotometer fitted with an Applied Photophysics RX pneumatic drive unit. For obtaining the macroscopic off-rate constant (Koff), two stock solutions were prepared as follows: a Ca^2+^-saturated indicator solution (30 mM MOPS, 1 mM CaCl2, 2 mM MgCl2, 100 mM KCl, ∼ 0.2-1 μM indicator, pH 7.2) and a BAPTA solution (30 mM MOPS, 100 mM KCl, 20 mM BAPTA, pH 7.2). The stopped-flow experiment was carried out at room temperature (∼23 °C) and the two solutions were mixed with an injection pressure of 3.5 bar. Excitation was set to 480 nm and emission was detected at 520 nm. The decay time (τ, s) was determined by fitting a double-exponential curve to the fluorescence response using Prism.

Macroscopic on-rate kinetics (Kobs) were obtained both for CaNeon and mCyRFP1- CaNeon by mixing the Ca^2+^-free buffer containing the protein (30 mM MOPS, 100 mM KCl, 1 mM MgCl2, 10 mM EGTA, ∼ 0.2-1 μM indicator, pH 7.2) and solutions containing increasing concentrations of CaCl2 (30 mM MOPS, 100 mM KCl, 8-20 mM CaCl2, 1 mM MgCl2, 10 mM EGTA, ∼ 0.2-1 μM indicator, pH 7.2). Concentrations of free Ca^2+^ were calculated using WEBMAXC STANDARD.

For measuring the pKa of the sensors a series of MOPS/MES (2-N-morpholino-ethane sulfonic acid) buffered solutions, supplemented with 1 mM Ca^2+^, were prepared. The pH values were adjusted in 0.5 pH steps from pH 5.5 to pH 8.5 using NaOH and HCl. In a bottom 96 well plate, triplicates of 200 μl of buffer containing 0.5-1 μM of protein were prepared for each pH value and all emission spectra were recorded. The relative fluorescence values at the emission maximum were plotted against the pH values and a sigmoidal fit was applied.

### Creation of the viral vector carrying mCyRFP1-CaNeon

We used the LV.Twitch-2B.miR-9.T construct containing the cytomegalovirus (CMV) promoter and four microRNA-9 target sequences (Brawek et al., 2017) as the parental vector to produce the LV. mCyRFP1-CaNeon.miR-9.T. We replaced Twitch-2B with mCyRFP1-CaNeon using a homologous recombination-based assembly molecular cloning approach. For this purpose, we propagated mCyRFP1-CaNeon by PCR amplification from a donor vector using primers that contain both mCyRFP1-CaNeon specific sequences and the flanking regions (15 nucleotides in length) located exactly up- and down-stream of the Twitch-2B sequence in the parenteral vector (forward primer: 5’-CTCTACTAGAGGATCCGCCACCATGGTGAGCAAGGGC-3’, reverse primer: 5’-GAGGTTGATTGTCGACTCAGTGGTATTTGTGAGCCAGGG-3’). The PCR product was digested with Dpn I, affinity-purified, and reserved for assembly molecular cloning. Then, the parental vector LV.Twitch-2B.miR-9.T was digested with the restriction enzymes BamH I and Sal I to remove the Twitch-2B sequence. The homologous recombination reaction was performed at a 1:2 ratio with 200 ng of the digested parenteral vector using the NEBuilder® HiFi DNA Assembly Cloning Kit from New England BioLabs.

### MicroRNA-9-regulated lentiviral vectors

MicroRNA-9 (miR-9) is a microRNA that promotes the degradation of mRNA with a specific complementary sequence. Because in mice miR-9 is expressed in virtually all CNS cells except microglia, it can be used for cell type-specific labeling of microglia (Akerblom et al., 2013; Brawek et al., 2017; Olmedillas et al., 2023). This cell type specificity, however, relies on the interaction between the endogenous miR-9 and the exogenous substrate and thus can be overridden, if the concentration of exogenous mRNA is high or the concentration of endogenous miR-9 is low.

Production of miR9-regulated lentiviral vectors was described previously (Brawek et al., 2017; Olmedillas et al., 2023). In brief, cell-free supernatants containing viral particles were produced by transient transfection of HEK293T packaging cells with the respective lentiviral construct along with the packing plasmids psPAX2 and pMD2G, following standard procedures. After 48 h the virus-containing culture supernatant was collected, filtered through a 0.45 µm pore-sized filter to remove cell debris and concentrated by centrifugation at 100000 g for 2 h at 4°C (Thermofisher WX Ultra80 centrifuge, Waltham, MA, USA). Pellets were resuspended in sterile PBS and stored at -80°C.

Four different miR-9-regulated lentiviral constructs were used in this study. Constructs containing the PGK promotor (Fig. 1B) enabled the expression of 3 different fluorophores (mCherry, Venus, or mTurquoise2) and the stochastic combination thereof (Olmedillas et al., 2023). The LV.mCyRFP1-CaNeon.miR-9.T construct contained a CMV promotor (Fig. 2C) and was used to label human microglia with a ratiometric Ca^2+^ indicator mCyRFP1-CaNeon. For this study, viral particle aliquots with no less than 10^8^ colony-forming units/ml were used.

### Two-photon imaging

Two-photon imaging was performed using a laser scanning microscope (Olympus Fluoview 1000 or Olympus Fluoview 300, Olympus, Tokyo, Japan) equipped with a 40x water-immersion objective (0.80 NA, Nikon, Tokyo, Japan) and coupled to a tunable titanium-sapphire laser (690-1040 nm excitation wavelength; MaiTai DeepSee, Spectra Physics, Santa Clara, CA, USA). During imaging the human slices were continuously perfused with Ringer’s solution (125 mM NaCl, 4.5 mM KCl, 26 mM NaHCO3, 1.25 mM NaH2PO4, 2.5 mM CaCl2, 1 mM MgCl2, and 20 mM glucose) at 32 °C and pH 7.4, when bubbled with 95% O2 and 5% CO2.

mTurquoise and mCherry were excited at 800 nm and mVenus at 990 nm. The emitted light was split by a 570 nm dichroic mirror and filtered with SP 570 nm or BP 630/92 nm, respectively. mCyRFP1-CaNeon was excited at 930 nm and a 570 nm dichroic mirror was utilized to split the emitted light into two separate channels for CaNeon (SP 570 nm) and mCyRFP1 (BP 630/92 nm), respectively. Microglial Ca^2+^ transients were recorded in 4D for 15 minutes with a resolution of 0.31 μm/pixel in the XY plane, Z step size of 2 µm and a sampling rate of 2 μs/pixel (0.3 to 0.5 Hz). 3D stacks were acquired at a spatial resolution of 0.31 μm/pixel and a Z step size of 1-2 μm.

### Ca^2+^ signal detection and analyses

Ca^2+^ signals were detected using the active voxels detection algorithm of the MATLAB (MathWorks) Begonia framework (Bjornstad et al., 2021), which permits the detection of pixel-by-pixel changes in fluorescence intensity. Within this framework, the fluorescence value *F* of each pixel is converted into a binary time series, where pixels with values other than 0 represent events. These events correspond to pixel grayscale values that exceed a user-determined threshold τ_*i*_, which is a function of the baseline grayscale values and the standard deviation of noise:

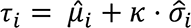

where *κ* is an empirically obtained coefficient that determines the height of the threshold. Depending on the image quality and the fluorescence intensity of CaNeon, *κ* values between 4 and 5 were chosen. μ_*i*_ is the baseline value of fluorescence for each pixel, and σ_*i*_ is the standard deviation of noise (Bjornstad et al., 2021).

The binary matrix of active pixels was then exported from MATLAB and further analyses were performed using a Python routine (Harris et al., 2020). The matrix’s pixels were subsequently added along the time axis T. As the value of active pixels is 1 and the value of inactive pixels is 0, the resulting image consists of the non-zero pixels that were active at any given point during the recording. Next, the binary map was segmented using the 1-connectivity of pixels (pixels connected only perpendicularly), resulting in separate localized ROAs that did not overlap in space throughout the entire registration. The locations of microdomains corresponding to each ROA were determined semi-automatically by assessing the degree of overlap between the ROA’s area and the area of the soma. This was achieved by applying the

Otsu threshold (Otsu, 1979) to the mean image of the recorded cell and calculating the overlap area.

Then, for both CaNeon and mCyRFP1 fluorescence, a mean intensity value for every frame in the area corresponding to each previously obtained ROA was calculated. The background-subtracted fluorescence ratio and the relative ratio change (ΔR/R) were calculated as follows:

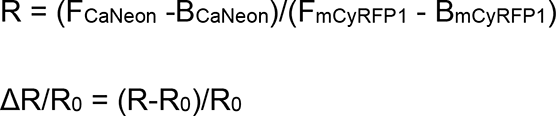

where the background fluorescence (B) for both channels was calculated as the mean intensity of a 10 x 10 pixels region located outside the analyzed cell and R0 is the baseline value of R. The baseline was approximated with a third-order weighted polynomial function. The peaks of ΔR/R traces were detected by utilizing the Python scipy-signal module (Virtanen et al., 2020). Only those peaks that exceeded the 4 x σnoise threshold were considered.

### Immunohistochemistry

After imaging experiments, human organotypic slices were fixed with 4% formaldehyde in PBS for 2 h at 4°C, washed in PBS and permeabilized using 0.25% Triton X in PBS for 15 min. Antibody staining was performed with free-floating slices at room temperature. Slices were treated with a blocking solution (5% normal goat serum, 3% bovine serum albumin (BSA) and 1% Triton X-100 in PBS) for one hour and incubated overnight with the primary antibodies (rabbit-anti-Iba-1, 1:200; Wako, USA; rat-anti-mCherry, 1:2000, Thermo Fisher Scientific). After washing in PBS, the slices were incubated with Alexa Fluor (AF) 488- and AF-594-conjugated secondary antibodies (1:1000, Invitrogen, Waltham, MA, USA) for 2 h in darkness; and later mounted on fluorescence-free Superfrost Plus microscope slides (Langenbrinck, Emmendingen, Germany) with Vectashield Mounting Medium (Vector Laboratories, Burlingame, CA, USA). Freshly resected cortical samples (Table 2) were fixed in 4% paraformaldehyde for 1 h at 4 °C and subsequently washed 3x15 min in PBS. The slices were transferred to a 15 % sucrose solution for 90 minutes and then incubated in a 30 % sucrose solution overnight. After washing the slices 3x15 min in PBS, they were incubated for 2 days at 4°C with the rabbit-anti-Iba-1 antibody (1:1000, Wako, USA). After washing with PBS, the slices were incubated for 24 hours at 4°C with Alexa Fluor 488 secondary antibody (1:750, Thermo Fisher Scientific). Finally, the slices were washed with PBS and mounted using Fluoromount-G (Thermo Fisher Scientific).

For immunolabeling of cortical slices of NG2-DsRed transgenic mice (Tg(Cspg4- DsRed.T1)1Akik/J), we used the brain tissue from our biobank, which was previously isolated and fixed with 4% formaldehyde in PBS for 2 h at 4°C, cryoprotected in 25% sucrose in PBS overnight at 4°C, embedded in Tissue Tek (Sakura Finetek, Torrance, CA, USA) and stored at -80°C. Antibody staining was performed with free-floating 50- μm-thick coronal sections at room temperature. The sections were rinsed in PBS, blocked in 1% bovine serum albumin (BSA) containing 0.3% Triton X-100 in PBS for 1 h at room temperature, followed by incubation with primary antibodies (rabbit-anti RFP, Rockland, diluted 1:1500 and goat anti-PDGFRα, R&D Systems, diluted 1:500) in 1% BSA, 0.1% Triton X-100 in PBS overnight. After washing in PBS, the slices were incubated with Alexa Fluor (AF) 488- and AF-594-conjugated secondary antibodies (1:1000, Invitrogen, Waltham, MA, USA) for 2 h in darkness; and later mounted on fluorescence-free Superfrost Plus microscope slides (Langenbrinck, Emmendingen, Germany) with Vectashield Mounting Medium (Vector Laboratories, Burlingame, CA, USA).

### Analyses of the soma size

For the reconstruction of microglial somata, images were processed using Imaris software (version 10.1.1, Bitplane, Oxford Instruments). Cell bodies were manually identified and the surface creation feature with manual adjustment of thresholds was used to delineate the boundaries of microglial somata. Morphological and intensity segmentation features were employed to exclude adjacent extracellular components (e.g. lipofuscin grains). Cells were only included in the analyses if a clear separation from the surrounding extracellular material was possible. Following reconstruction, several parameters including 3 bounding box dimensions (i.e. cell diameters in X, Y, Z), the surface area and volume as well as sphericity were extracted from Imaris software.

### Statistical analyses and data presentation

Statistical tests were performed in Python using the Scipy library. The one-sample Shapiro-Wilk test was used to check for the normality of the data distribution and the Levene’s test was used to test for homoscedasticity. Comparisons of more than two independent variables were performed using the Kruskal-Wallis test followed by the Holm-Bonferroni post hoc test for multiple comparisons. All statistical tests were two-sided. The P values ≤ 0.05 were considered significant. If not otherwise indicated, data is presented as median ± interquartile range. Lines of boxes in box plots represent 25th and 75th, and whiskers 10th and 90th percentiles.

## Supplementary figures

Figure S1. Spectroscopic *in vitro* properties of CaNeon and mCyRFP1-CaNeon.

(A) The amino acid sequence of CaNeon: mNeonGreen (green), TnCmin (orange), linkers (black). The amino acid changes introduced during sensor optimization and thus differing from parental NRS1.2 are underlined. (**B-C**) Emission spectra of CaNeon

(B) and mCyRFP1-CaNeon (**C**) at increasing concentrations of Ca^2+^ (from light to dark green: 0, 0.027, 0.065, 0.1, 0.225, 0.361, 0.602, 0.853, 1.73, 2.85, 7.37 and 14.9 mM).

(**D**) Summary of the main spectroscopic parameters of both indicators (see Materials and methods for measurement details). (**E-H**) Ca^2+^ responsiveness of mCyRFP1- CaNeon in HEK 293 cells, cultured using the standard protocol (Loew et al., 2010) and transduced with miR9-regulated lentiviral vectors. Average intensity images of a sample FOV recorded in green (**E**, left) and red (**E**, right) channels, and their overlays (**F**), taken at different time points (t1 and t2, see also (**G-H**)) during the measurement, illustrate the baseline level of CaNeon and mCyRFP1 fluorescence (**E** and **F**, left) and fluorescence changes (**F**, right) caused by a 1-min-long bath application of 20 mM caffeine, dissolved in the Ringer’s solution. The ΔF/F and ΔR/R signals recorded in regions of interest delineated in (**F**, left) are shown in (**G**) and (**H**), respectively.

Figure S2. **miR-9 regulated lentiviral vectors weekly label a non-microglial population of human cells.** (**A**) MIP (4-16 µm depth) image of a fixed human organotypic brain slice, labeled with antibodies against a microglia/macrophage marker Iba-1 (magenta) and mCyRFP1-CaNeon (green). (**B**) MIP (13-21 µm depth) image of a fixed cortical slice of a mouse mutant expressing DsRed under the NG2 promotor, thus labeling NG2-positive oligodendrocyte precursor cells and pericytes (Zhu et al., 2008). The tissue was labeled with antibodies against RFP (green, recognizing DsRed) and NG2 cell marker PDGFRα (magenta). Note the similarity in morphology between the weakly-labeled non-microglial human cells (**A**, arrows) and mouse pericytes (**B**, arrows).

**Figure S3. Morphology of microglia from freshly resected human tissue.** (**A**) Sample MIP images of ramified, hypertrophic and ameboid microglia (1-13, 31-37, 34- 39 µm depth, respectively), labeled with antibodies against Iba-1. Yellow-reddish structures likely represent the autofluorescent wear-and-tear pigment lipofuscin, known to have broad excitation and emission spectra (Eichhoff et al., 2008). (**B**-**E**) Plots, illustrating the distributions of the mean diameter (**B**), surface area (**C**), volume (**D**) and sphericity (**E**) of microglial somata from freshly resected (n=175 cells; black) and cultured (n=55 cells, red) human tissue. Vertical broken lines indicate the maximal value of the respective parameter in freshly resected microglia. Two-sample Kolmogorov-Smirnov test, P<10^-4^ (**B-D**) and P=0.64 (**E**).

**Figure S4. Characteristics of morphotype-specific microglial Ca^2+^ transients in different subcellular compartments.** (**A-C**) Box plots showing the amplitude (**A**; P=6.5*10^-10^, 4.8*10^-14^ and 5.2*10^-3^ for comparison of processes to soma and processes, processes to soma and soma and processes to soma, respectively, in ramified microglia (here and below Kruskal-Wallis test followed by Holm-Bonferroni post hoc test for multiple comparisons)), FWHM (**B**; P=4*10^-3^ for comparison of microglial processes to soma), and AUC (**C**; P=8.7*10^-9^, 7.4*10^-10^ and 3.4*10^-4^ for comparison of processes to soma and processes, processes to soma and soma and processes to soma, respectively, in ramified microglia) of Ca^2+^ transients recorded in different subcellular compartments of ramified microglia. (**D-F**) Box plots showing themplitude (**D**; P=2.4*10^-8^, 9.2*10^-6^ and 0.05 for comparison of processes to soma and processes, processes to soma and soma and processes to soma, respectively), FWHM (**E**), and AUC (**F**; P=1.4*10^-9^, 2.4*10^-6^ and 0.02 for comparison of processes to soma and processes, processes to soma and soma and processes to soma, respectively) of Ca^2+^ transients recorded in different subcellular compartments of hypertrophic microglia. (**G-I**) Box plots showing the amplitude (**G**; P=2.8*10^-3^), FWHM (**H**; P=1.4*10^-3^), and AUC (**I**; P=1.1*10^-4^) of Ca^2+^ transients recorded in soma and processes as well as soma of amoeboid microglia.

**Movie S1. Process motility of human microglia.** An overlay of CaNeon (green) and mCyRFP1 (red) channels, showing a microglial cell, vividly moving its processes. Each channel is an average intensity projection of 6 images (10-16 µm, step 1 µm). Note several increases in green fluorescence (Ca^2+^ transients) accompanying process movement. Scale bar: 10 µm. Frame rate: 4.57 s, the movie plays 100 times faster.

**Movie S2. Localized Ca^2+^ signaling in phagocytic cups.** An overlay of CaNeon (green) and mCyRFP1 (red) channels, showing a microglial cell with several phagocytic cups. Each channel is an average intensity projection of 9 images (16-24 µm, step 1 µm). Note several increases in green fluorescence (Ca^2+^ transients) localized in the cup vicinity (arrowheads) and one cup, which is pulled towards the parent process (arrow). Scale bar: 10 µm. Frame rate: 4.94 s, the movie plays 100 times faster.

## References

1. Abels, E.R., L. Nieland, S. Hickman, M.L.D. Broekman, J. El Khoury, and S.L.N. Maas. 2021. Comparative analysis identifies similarities between the human and murine microglial sensomes. Int J Mol Sci 22:

2. Akerblom, M., R. Sachdeva, L. Quintino, E.E. Wettergren, K.Z. Chapman, G. Manfre, O. Lindvall, C. Lundberg, and J. Jakobsson. 2013. Visualization and genetic modification of resident brain microglia using lentiviral vectors regulated by microRNA-9. Nat Commun 4:1770.

3. Askew, K., K. Li, A. Olmos-Alonso, F. Garcia-Moreno, Y. Liang, P. Richardson, T. Tipton, M.A. Chapman, K. Riecken, S. Beccari, A. Sierra, Z. Molnar, M.S. Cragg, O. Garaschuk, V.H. Perry, and D. Gomez-Nicola. 2017. Coupled Proliferation and Apoptosis Maintain the Rapid Turnover of Microglia in the Adult Brain. Cell Rep 18:391–405.

4. Bjornstad, D.M., K.S. Abjorsbraten, E. Hennestad, C. Cunen, G.H. Hermansen, L. Bojarskaite, K.H. Pettersen, K. Vervaeke, and R. Enger. 2021. Begonia-A Two-Photon Imaging Analysis Pipeline for Astrocytic Ca(2+) Signals. Front Cell Neurosci 15:681066.

5. Bolmont, T., F. Haiss, D. Eicke, R. Radde, C.A. Mathis, W.E. Klunk, S. Kohsaka, M. Jucker, and M.E. Calhoun. 2008. Dynamics of the microglial/amyloid interaction indicate a role in plaque maintenance. J Neurosci 28:4283–4292.

6. Brawek, B., and O. Garaschuk. 2017. Monitoring in vivo function of cortical microglia. Cell Calcium 64:109–117.

7. Brawek, B., Y. Liang, D. Savitska, K. Li, N. Fomin-Thunemann, Y. Kovalchuk, E. Zirdum, J. Jakobsson, and O. Garaschuk. 2017. A new approach for ratiometric in vivo calcium imaging of microglia. Sci Rep 7:6030.

8. Chen, T.W., T.J. Wardill, Y. Sun, S.R. Pulver, S.L. Renninger, A. Baohan, E.R. Schreiter, R.A. Kerr, M.B. Orger, V. Jayaraman, L.L. Looger, K. Svoboda, and D.S. Kim. 2013. Ultrasensitive fluorescent proteins for imaging neuronal activity. Nature 499:295–300.

9. Dolmetsch, R.E., K. Xu, and R.S. Lewis. 1998. Calcium oscillations increase the efficiency and specificity of gene expression. Nature 392:933–936.

10. Eichhoff, G., B. Brawek, and O. Garaschuk. 2011. Microglial calcium signal acts as a rapid sensor of single neuron damage in vivo. Biochim Biophys Acta 1813:1014–1024.

11. Eichhoff, G., M.A. Busche, and O. Garaschuk. 2008. In vivo calcium imaging of the aging and diseased brain. Eur J Nucl Med Mol Imaging 35 Suppl 1:S99–106.

12. Franco Bocanegra, D.K., J.A.R. Nicoll, and D. Boche. 2018. Innate immunity in Alzheimer’s disease: the relevance of animal models? J Neural Transm (Vienna) 125:827–846.

13. Galatro, T.F., I.R. Holtman, A.M. Lerario, I.D. Vainchtein, N. Brouwer, P.R. Sola, M.M. Veras, T.F. Pereira, R.E.P. Leite, T. Moller, P.D. Wes, M.C. Sogayar, J.D. Laman, W. den Dunnen, C.A. Pasqualucci, S.M. Oba-Shinjo, E. Boddeke, S.K.N. Marie, and B.J.L. Eggen. 2017. Transcriptomic analysis of purified human cortical microglia reveals age-associated changes. Nat Neurosci 20:1162–1171.

14. Garaschuk, O., and A. Verkhratsky. 2019. Physiology of Microglia. Methods Mol Biol 2034:27–40.

15. Garaschuk, O., Y. Yaari, and A. Konnerth. 1997. Release and sequestration of calcium by ryanodine-sensitive stores in rat hippocampal neurones. J Physiol 502 (Pt 1):13–30.

16. Geirsdottir, L., E. David, H. Keren-Shaul, A. Weiner, S.C. Bohlen, J. Neuber, A. Balic, A. Giladi, F. Sheban, C.A. Dutertre, C. Pfeifle, F. Peri, A. Raffo-Romero, J. Vizioli, K. Matiasek, C. Scheiwe, S. Meckel, K. Matz-Rensing, F. van der Meer, F.R. Thormodsson, C. Stadelmann, N. Zilkha, T. Kimchi, F. Ginhoux, I. Ulitsky, D. Erny, I. Amit, and M. Prinz. 2019. Cross-Species Single-Cell Analysis Reveals Divergence of the Primate Microglia Program. Cell 179:1609–1622 e1616.

17. Ginhoux, F., M. Greter, M. Leboeuf, S. Nandi, P. See, S. Gokhan, M.F. Mehler, S.J. Conway, L.G. Ng, E.R. Stanley, I.M. Samokhvalov, and M. Merad. 2010. Fate mapping analysis reveals that adult microglia derive from primitive macrophages. Science 330:841–845.

18. Gosselin, D., D. Skola, N.G. Coufal, I.R. Holtman, J.C.M. Schlachetzki, E. Sajti, B.N. Jaeger, C. O’Connor, C. Fitzpatrick, M.P. Pasillas, M. Pena, A. Adair, D.D. Gonda, M.L. Levy, R.M. Ransohoff, F.H. Gage, and C.K. Glass. 2017. An environment-dependent transcriptional network specifies human microglia identity. Science 356:

19. Granzotto, A., A. McQuade, J.P. Chadarevian, H. Davtyan, S.L. Sensi, I. Parker, M. Blurton-Jones, and I. Smith. 2024. ER and SOCE Ca^2+^ signals are not required for directed cell migration in human microglia. *bioRxiv*

20. Hanisch, U.K., and H. Kettenmann. 2007. Microglia: active sensor and versatile effector cells in the normal and pathologic brain. Nat Neurosci 10:1387–1394.

21. Harris, C.R., K.J. Millman, S.J. van der Walt, R. Gommers, P. Virtanen, D. Cournapeau, E. Wieser, J. Taylor, S. Berg, N.J. Smith, R. Kern, M. Picus, S. Hoyer, M.H. van Kerkwijk, M. Brett, A. Haldane, J.F. Del Rio, M. Wiebe, P. Peterson, P. Gerard-Marchant, K. Sheppard, T. Reddy, W. Weckesser, H. Abbasi, C. Gohlke, and T.E. Oliphant. 2020. Array programming with NumPy. Nature 585:357–362.

22. Hickman, S.E., N.D. Kingery, T.K. Ohsumi, M.L. Borowsky, L.C. Wang, T.K. Means, and J. El Khoury. 2013. The microglial sensome revealed by direct RNA sequencing. Nat Neurosci 16:1896–1905.

23. Jairaman, A., A. McQuade, A. Granzotto, Y.J. Kang, J.P. Chadarevian, S. Gandhi, I. Parker, I. Smith, H. Cho, S.L. Sensi, S. Othy, M. Blurton-Jones, and M.D. Cahalan. 2022. TREM2 regulates purinergic receptor-mediated calcium signaling and motility in human iPSC-derived microglia. Elife 11:

24. Janks, L., C.V.R. Sharma, and T.M. Egan. 2018. A central role for P2X7 receptors in human microglia. J Neuroinflammation 15:325.

25. Jantti, H., V. Sitnikova, Y. Ishchenko, A. Shakirzyanova, L. Giudice, I.F. Ugidos, M. Gomez-Budia, N. Korvenlaita, S. Ohtonen, I. Belaya, F. Fazaludeen, N. Mikhailov, M. Gotkiewicz, K. Ketola, S. Lehtonen, J. Koistinaho, K.M. Kanninen, D. Hernandez, A. Pebay, R. Giugno, P. Korhonen, R. Giniatullin, and T. Malm. 2022. Microglial amyloid beta clearance is driven by PIEZO1 channels. J Neuroinflammation 19:147.

26. Keren-Shaul, H., A. Spinrad, A. Weiner, O. Matcovitch-Natan, R. Dvir-Szternfeld, T.K. Ulland, E. David, K. Baruch, D. Lara-Astaiso, B. Toth, S. Itzkovitz, M. Colonna, M. Schwartz, and I. Amit. 2017. A Unique Microglia Type Associated with Restricting Development of Alzheimer’s Disease. Cell 169:1276–1290 e1217.

27. Konttinen, H., M.E.C. Cabral-da-Silva, S. Ohtonen, S. Wojciechowski, A. Shakirzyanova, S. Caligola, R. Giugno, Y. Ishchenko, D. Hernandez, M.F. Fazaludeen, S. Eamen, M.G. Budia, I. Fagerlund, F. Scoyni, P. Korhonen, N. Huber, A. Haapasalo, A.W. Hewitt, J. Vickers, G.C. Smith, M. Oksanen, C. Graff, K.M. Kanninen, S. Lehtonen, N. Propson, M.P. Schwartz, A. Pebay, J. Koistinaho, L. Ooi, and T. Malm. 2019. PSEN1DeltaE9, APPswe, and APOE4 Confer Disparate Phenotypes in Human iPSC-Derived Microglia. Stem Cell Reports 13:669-683.

29. Laviv, T., B.B. Kim, J. Chu, A.J. Lam, M.Z. Lin, and R. Yasuda. 2016. Simultaneous dual-color fluorescence lifetime imaging with novel red-shifted fluorescent proteins. Nat Methods 13:989–992.

30. Loew, R., Y. Meyer, K. Kuehlcke, L. Gama-Norton, D. Wirth, H. Hauser, S. Stein, M. Grez, S. Thornhill, A. Thrasher, C. Baum, and A. Schambach. 2010. A new PG13-based packaging cell line for stable production of clinical-grade self-inactivating gamma-retroviral vectors using targeted integration. Gene Ther 17:272–280.

31. Logiacco, F., P. Xia, S.V. Georgiev, C. Franconi, Y.J. Chang, B. Ugursu, A. Sporbert, R. Kuhn, H. Kettenmann, and M. Semtner. 2021. Microglia sense neuronal activity via GABA in the early postnatal hippocampus. Cell Rep 37:110128.

32. Lushchak, V.I., M. Duszenko, D.V. Gospodaryov, and O. Garaschuk. 2021. Oxidative Stress and Energy Metabolism in the Brain: Midlife as a Turning Point. Antioxidants (Basel) 10:

33. Ma, C., B. Li, D. Silverman, X. Ding, A. Li, C. Xiao, G. Huang, K. Worden, S. Muroy, W. Chen, Z. Xu, C.F. Tso, Y. Huang, Y. Zhang, Q. Luo, K. Saijo, and Y. Dan. 2024. Microglia regulate sleep through calcium-dependent modulation of norepinephrine transmission. Nat Neurosci 249–258.

34. Mancuso, R., J. Van Den Daele, N. Fattorelli, L. Wolfs, S. Balusu, O. Burton, A. Liston, A. Sierksma, Y. Fourne, S. Poovathingal, A. Arranz-Mendiguren, C. Sala Frigerio, C. Claes, L. Serneels, T. Theys, V.H. Perry, C. Verfaillie, M. Fiers, and B. De Strooper. 2019. Stem-cell-derived human microglia transplanted in mouse brain to study human disease. Nat Neurosci 22:2111–2116.

35. McGeachan, R.I., S. Meftah, L.W. Taylor, J.H. Catterson, D. Negro, J. Tulloch, J.L. Rose, F. Gobbo, I. Liaquat, T.L. Spires-Jones, S.A. Booker, P.M. Brennan, and C.S. Durrant. 2024. Opposing roles of physiological and pathological amyloid-β on synapses in live human brain slice cultures. bioRxiv 2024.2002.2016.580676.

36. Mondo, E., S.C. Becker, A.G. Kautzman, M. Schifferer, C.E. Baer, J. Chen, E.J. Huang, M. Simons, and D.P. Schafer. 2020. A developmental analysis of juxtavascular microglia dynamics and interactions with the vasculature. J Neurosci 40:6503–6521.

37. Nimmerjahn, A., F. Kirchhoff, and F. Helmchen. 2005. Resting microglial cells are highly dynamic surveillants of brain parenchyma in vivo. Science 308:1314–1318.

38. Olmedillas del Moral, M.O., N. Fröhlich, K. Figarella, N. Mojtahedi, and O. Garaschuk. 2020. Effect of caloric restriction on the functional properties of aging microglia. Front Immunol 11:750.

39. Olmedillas, M., B. Brawek, K. Li, C. Richter, and O. Garaschuk. 2023. Plaque vicinity as a hotspot of microglial turnover in a mouse model of Alzheimer’s disease. Glia 71:2884–2901.

40. Otsu, N. 1979. A threshold selection method from gray-level histograms. IEEE Transactions on systems, man, and cybernetics SMC-9:62–66.

41. Pan, K., and O. Garaschuk. 2022. The role of intracellular calcium-store-mediated calcium signals in in vivo sensor and effector functions of microglia. J Physiol 4203–4215.

42. Pozner, A., B. Xu, S. Palumbos, J.M. Gee, P. Tvrdik, and M.R. Capecchi. 2015. Intracellular calcium dynamics in cortical microglia responding to focal laser injury in the PC::G5-tdT reporter mouse. Front Mol Neurosci 8:12.

43. Que, Z., M.I. Olivero-Acosta, I. Chen, J. Zhang, K. Wettschurack, J. Wu, T. Xiao, C.M. Otterbacher, M. Wang, H. Harlow, N. Cui, X. Chen, B. Deming, M. Halurkar, Y. Zhao, J.C. Rochet, R. Xu, A.L. Brewster, L.J. Wu, C. Yuan, W.C. Skarnes, and Y. Yang. 2023. Human iPSC-derived microglia sense and dampen hyperexcitability of cortical neurons carrying the epilepsy-associated SCN2A- L1342P mutation. *bioRxiv*

44. Reu, P., A. Khosravi, S. Bernard, J.E. Mold, M. Salehpour, K. Alkass, S. Perl, J. Tisdale, G. Possnert, H. Druid, and J. Frisen. 2017. The Lifespan and Turnover of Microglia in the Human Brain. Cell Rep 20:779–784.

45. Riester, K., B. Brawek, D. Savitska, N. Frohlich, E. Zirdum, N. Mojtahedi, M.T. Heneka, and O. Garaschuk. 2020. In vivo characterization of functional states of cortical microglia during peripheral inflammation. Brain Behav Immun 87:243–255.

46. Schafer, S.T., A.A. Mansour, J.C.M. Schlachetzki, M. Pena, S. Ghassemzadeh, L. Mitchell, A. Mar, D. Quang, S. Stumpf, I.S. Ortiz, A.J. Lana, C. Baek, R. Zaghal, C.K. Glass, A. Nimmerjahn, and F.H. Gage. 2023. An in vivo neuroimmune organoid model to study human microglia phenotypes. Cell 186:2111–2126 e2120.

47. Schwarz, N., U.B.S. Hedrich, H. Schwarz, A.H. P, N. Dammeier, E. Auffenberg, F. Bedogni, J.B. Honegger, H. Lerche, T.V. Wuttke, and H. Koch. 2017. Human Cerebrospinal fluid promotes long-term neuronal viability and network function in human neocortical organotypic brain slice cultures. Sci Rep 7:12249.

48. Schwarz, N., B. Uysal, M. Welzer, J.C. Bahr, N. Layer, H. Loffler, K. Stanaitis, H. Pa, Y.G. Weber, U.B. Hedrich, J.B. Honegger, A. Skodras, A.J. Becker, T.V. Wuttke, and H. Koch. 2019. Long-term adult human brain slice cultures as a model system to study human CNS circuitry and disease. Elife 8:

49. Seifert, S., M. Pannell, W. Uckert, K. Farber, and H. Kettenmann. 2011. Transmitter- and hormone-activated Ca^2+^ responses in adult microglia/brain macrophages in situ recorded after viral transduction of a recombinant Ca^2+^ sensor. Cell Calcium 49:365–375.

50. Shaner, N.C., G.G. Lambert, A. Chammas, Y. Ni, P.J. Cranfill, M.A. Baird, B.R. Sell, J.R. Allen, R.N. Day, M. Israelsson, M.W. Davidson, and J. Wang. 2013. A bright monomeric green fluorescent protein derived from Branchiostoma lanceolatum. Nat Methods 10:407–409.

51. Torres-Platas, S.G., S. Comeau, A. Rachalski, G.D. Bo, C. Cruceanu, G. Turecki, B. Giros, and N. Mechawar. 2014. Morphometric characterization of microglial phenotypes in human cerebral cortex. J Neuroinflamm 11:12.

52. Tsien, R.Y. 1989. Fluorescent indicators of ion concentrations. Methods Cell Biol 30:127–156.

53. Umpierre, A.D., L.L. Bystrom, Y. Ying, Y.U. Liu, G. Worrell, and L.J. Wu. 2020. Microglial calcium signaling is attuned to neuronal activity in awake mice. Elife 9:e56502.

54. Valiente-Gabioud, A.A., I. Garteizgogeascoa Suner, A. Idziak, A. Fabritius, J. Basquin, J. Angibaud, U.V. Nagerl, S.P. Singh, and O. Griesbeck. 2023. Fluorescent sensors for imaging of interstitial calcium. Nat Commun 14:6220.

55. Virtanen, P., R. Gommers, T.E. Oliphant, M. Haberland, T. Reddy, D. Cournapeau, E. Burovski, P. Peterson, W. Weckesser, J. Bright, S.J. van der Walt, M. Brett, J. Wilson, K.J. Millman, N. Mayorov, A.R.J. Nelson, E. Jones, R. Kern, E. Larson, C.J. Carey, I. Polat, Y. Feng, E.W. Moore, J. VanderPlas, D. Laxalde, J. Perktold, R. Cimrman, I. Henriksen, E.A. Quintero, C.R. Harris, A.M. Archibald, A.H. Ribeiro, F. Pedregosa, P. van Mulbregt, and C. SciPy. 2020. SciPy 1.0: fundamental algorithms for scientific computing in Python. Nat Methods 17:261–272.

56. Wang, F.S., W. Wang, S.M. Gu, D. Qi, N.A. Smith, W.G. Peng, W. Dong, J.J. Yuan, B.B. Zhao, Y. Mao, P. Cao, Q.R. Lu, L.A. Shapiro, S.S. Yi, E.X. Wu, and J.H. Huang. 2023. Distinct astrocytic modulatory roles in sensory transmission during sleep, wakefulness, and arousal states in freely moving mice. Nat Commun 14:2186.

57. Xia, P., F. Logiacco, Y. Huang, H. Kettenmann, and M. Semtner. 2021. Histamine triggers microglial responses indirectly via astrocytes and purinergic signaling. Glia 69:2291–2304.

58. Zhang, Y., U. Werling, and W. Edelmann. 2012. SLiCE: a novel bacterial cell extract-based DNA cloning method. Nucleic Acids Res 40:e55.

59. Zhou, Y., W.M. Song, P.S. Andhey, A. Swain, T. Levy, K.R. Miller, P.L. Poliani, M. Cominelli, S. Grover, S. Gilfillan, M. Cella, T.K. Ulland, K. Zaitsev, A. Miyashita, T. Ikeuchi, M. Sainouchi, A. Kakita, D.A. Bennett, J.A. Schneider, M.R. Nichols, S.A. Beausoleil, J.D. Ulrich, D.M. Holtzman, M.N. Artyomov, and M. Colonna. 2020. Human and mouse single-nucleus transcriptomics reveal TREM2- dependent and TREM2-independent cellular responses in Alzheimer’s disease. Nat Med 26:131–142.

60. Zhu, X.Q., D.E. Bergles, and A. Nishiyama. 2008. NG2 cells generate both oligodendrocytes and gray matter astrocytes. Development 135:145–157.

