## Supplemental Figures for "Morphotype-specific calcium signaling in human microglia"

**Movie S2. Localized $Ca^{2+}$ signaling in phagocytic cups.** An overlay of CaNeon (green) and mCyRFP1 (red) channels, showing a microglial cell with several phagocytic cups. Each channel is an average intensity projection of 9 images (16-24 µm, step 1 µm). Note several increases in green fluorescence ($Ca^{2+}$ transients) localized in the cup vicinity (arrowheads) and one cup, which is pulled towards the parent process (arrow). Scale bar: 10 µm. Frame rate: 4.94 s, the movie plays 100 times faster.

**A**

```
           10         20         30         40         50         60
MVSKGEEDNMGSLPATHELHIFGSINGVDFDMVGQGSGNPNDGYEELNLKSTKGDLQFSP
           70         80         90        100        110        120
WILVPHIGYGFHQYLPYPDGMSPFQAAMVDGSGYQVHRTMQFEDGASLTVNYRYTYEGSH
          130        140        150        160        170        180
IKGEAQVKGTGFPADGPVMTNSLTAADLGWDSEEELSEYFRIFDFDGNGFIDREEFGDII
          190        200        210        220        230        240
RLTGEQLTDEDVDEIFGGSDTDKNGRIDFDEFLKMVENVQLTDNNRSKKTYPNDKTIIST
          250        260        270        280        290        300
YKWSYTTDNGKRYRCTARTTYTFAKPMAANYLKNQPMYVFRKTELKHSKTELNFKEWQKAFTD
```

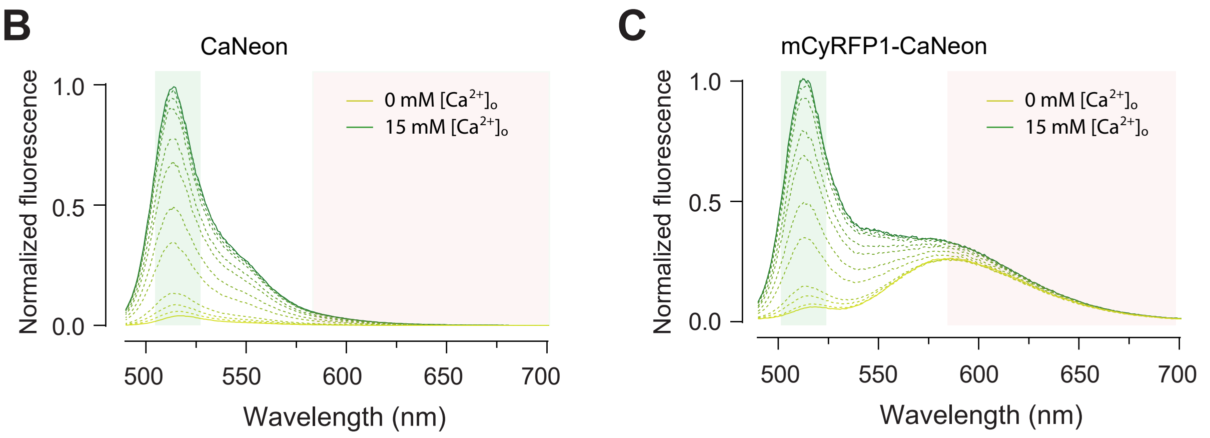


**D**

| | $\Delta F/F_0$ | $\Delta R/R_0$ | EC ($mM^{-1}cm^{-1}$) | QY | Brightness | $K^*_{obs}$ ($s^{-1}$) | Kd (nM) | Hill coeff. | $K_{off}$ ($s^{-1}$) | pKa |
|---|---|---|---|---|---|---|---|---|---|---|
| CaNeon | 30 | | 74 | 0.6 | 46 | 4.9 | 385 | 1.5 | 1.7 | 6.5 |
| mCyRFP1-CaNeon | | 16 | 70 | 0.6 | 43 | 4.2 | 401 | 1.6 | 1.7 | 6.6 |

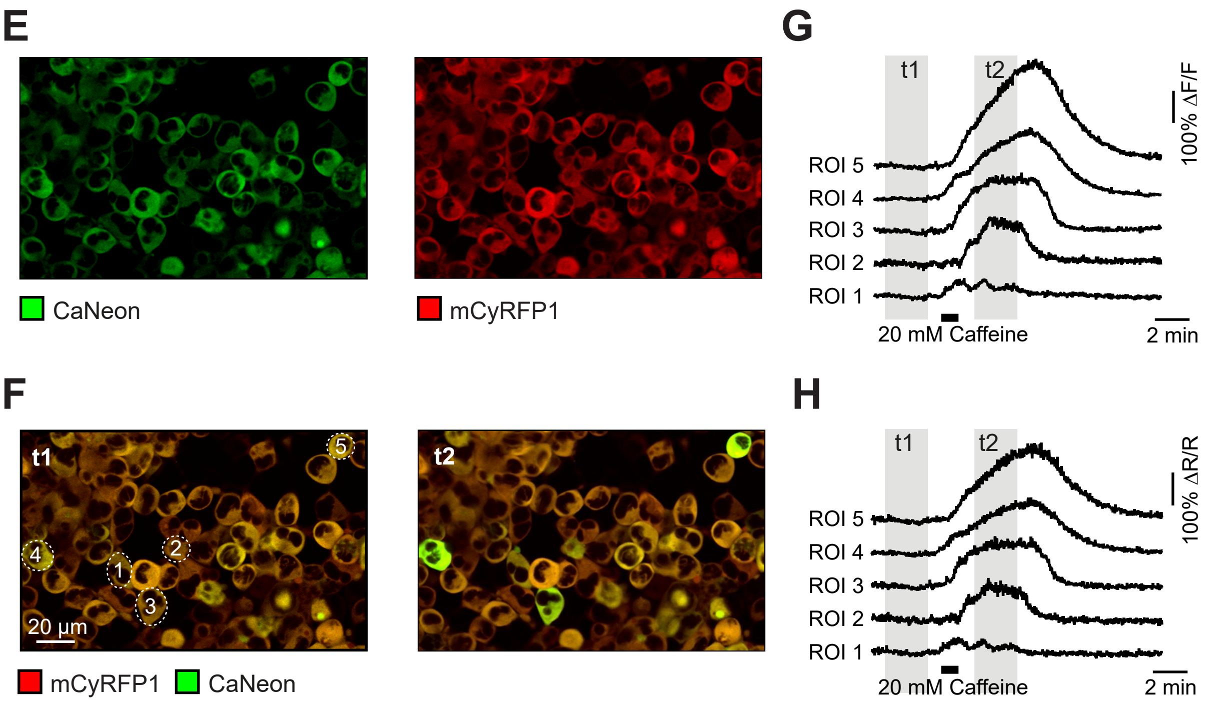

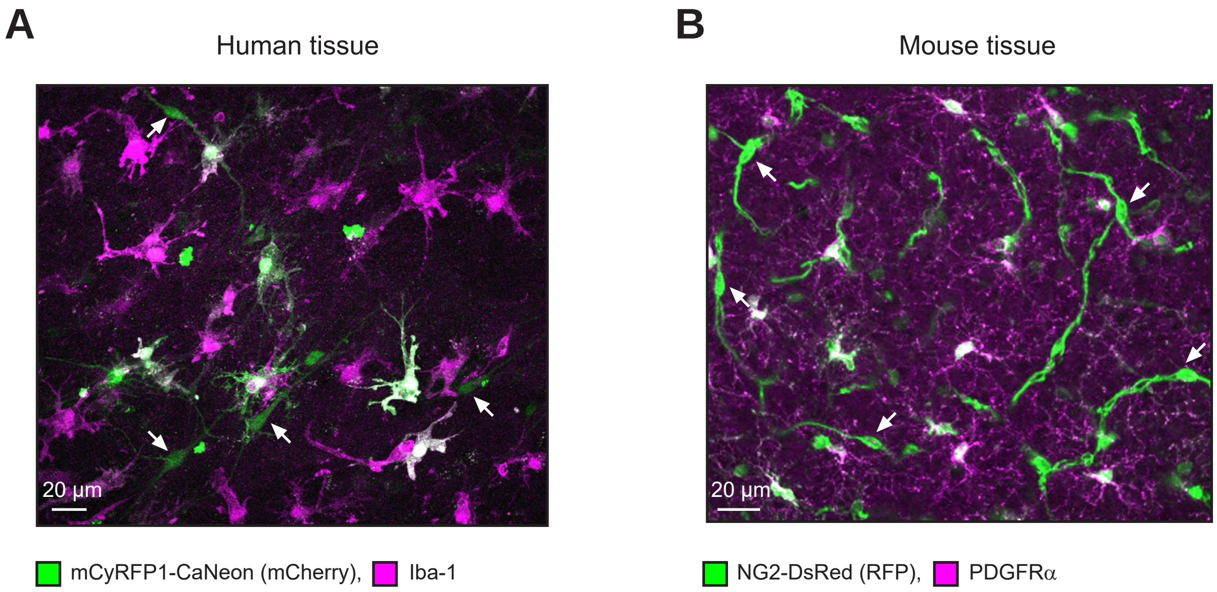

A
Human tissue
20 µm
mCyRFP1-CaNeon (mCherry),
Iba-1
B
Mouse tissue
20 µm
NG2-DsRed (RFP),
PDGFRα

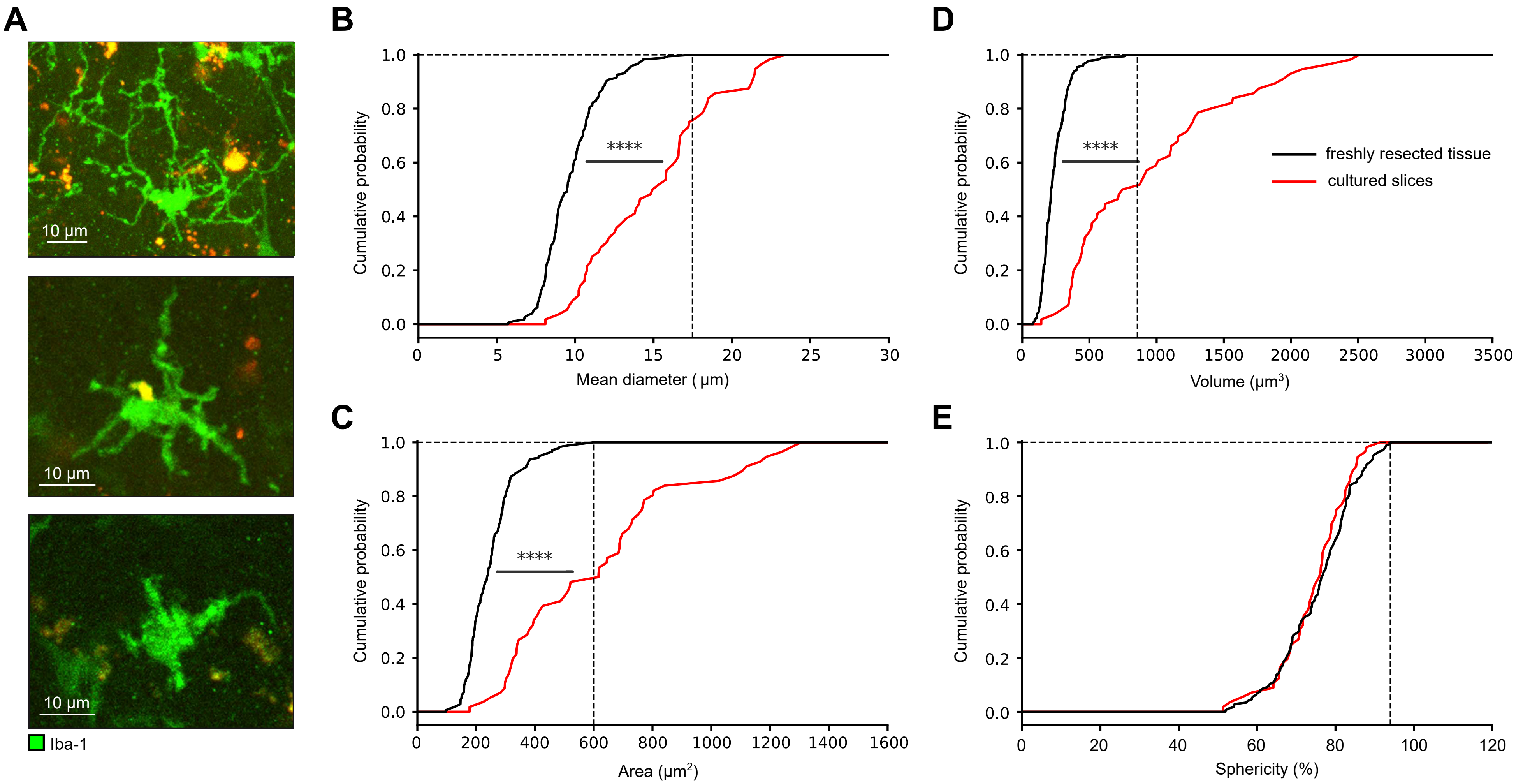

A
10 µm
10 µm
10 µm
Iba-1
B
****
Cumulative probability
Mean diameter ( µm)
C
****
Cumulative probability
Area (µm2)
D
****
Cumulative probability
Volume (µm3)
freshly resected tissue
cultured slices
E
Cumulative probability
Sphericity (%)

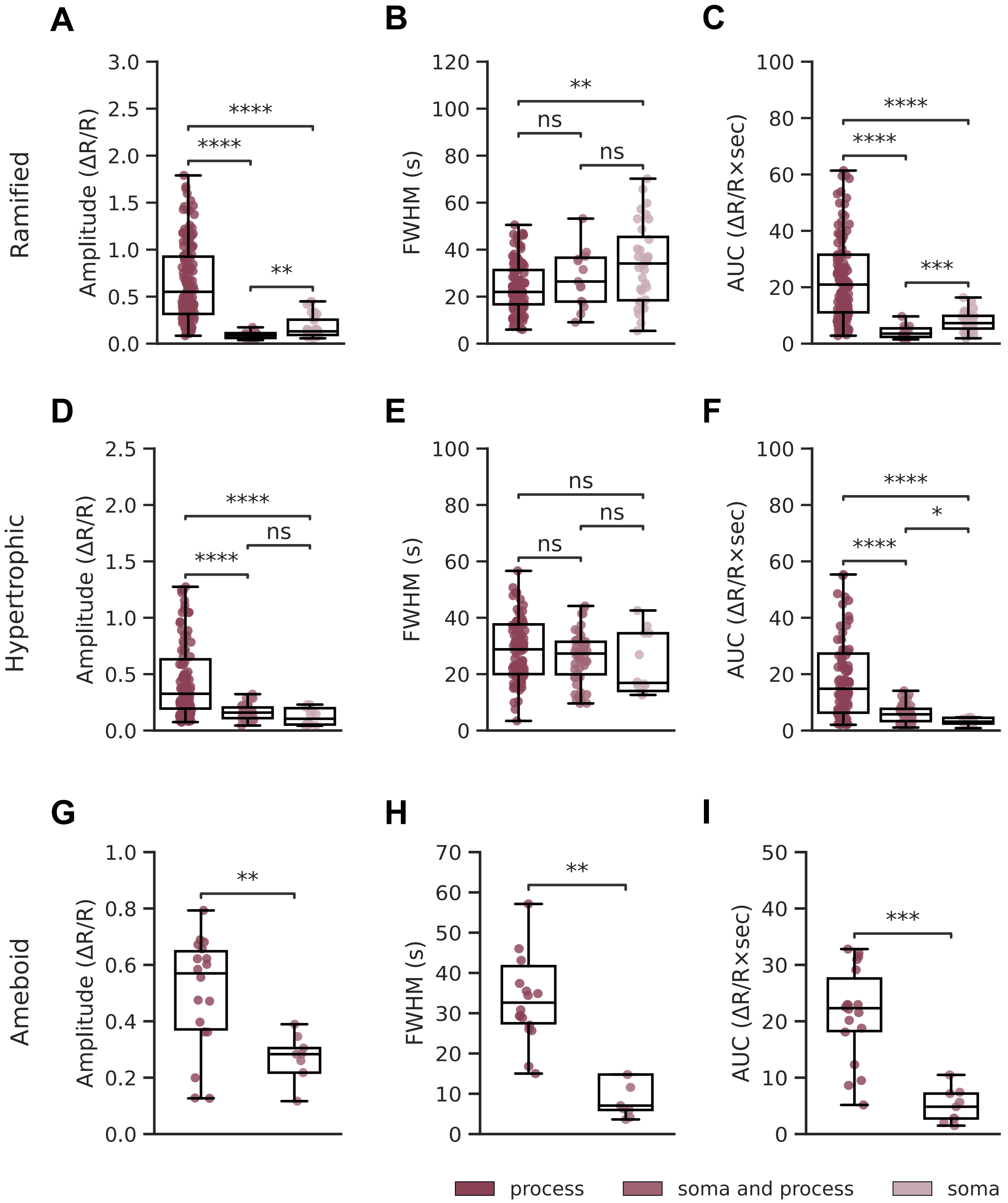
A
B
C
Ramified
Amplitude (ΔR/R)
FWHM (s)
AUC (ΔR/R×sec)
****
**
ns
***
D
E
F
Hypertrophic
*
G
H
I
Ameboid
process
soma and process
soma
